# Molecular Determinants and Signaling Effects of PKA RIα Phase Separation

**DOI:** 10.1101/2023.12.10.570836

**Authors:** Julia C. Hardy, Emily H. Pool, Jessica G.H. Bruystens, Xin Zhou, Qingrong Li, Daojia R. Zhou, Max Palay, Gerald Tan, Lisa Chen, Jaclyn L.C. Choi, Ha Neul Lee, Stefan Strack, Dong Wang, Susan S. Taylor, Sohum Mehta, Jin Zhang

**Affiliations:** Department of Pharmacology, University of California, San Diego, La Jolla, CA 92093, USA; Shu Chien-Gene Lay Department of Bioengineering, University of California, San Diego, La Jolla, CA 92093, USA; Department of Chemistry and Biochemistry, University of California, San Diego, La Jolla, CA 92093, USA; Skaggs School of Pharmacy and Pharmaceutical Sciences, University of California, San Diego, La Jolla, CA 92093, USA; Department of Pharmacology, University of Iowa, Iowa City, IA 52242, USA; Moores Cancer Center, University of California, San Diego, La Jolla, CA 92093, USA

**Keywords:** Biomolecular Condensates, Intrinsically Disordered Region, Protein Kinase, Signaling Compartmentation

## Abstract

Spatiotemporal regulation of intracellular signaling molecules, such as the 3’,5’-cyclic adenosine monophosphate (cAMP)-dependent protein kinase (PKA), ensures the specific execution of various cellular functions. Liquid-liquid phase separation (LLPS) of the ubiquitously expressed PKA regulatory subunit RIα was recently identified as a major driver of cAMP compartmentation and signaling specificity. However, the molecular determinants of RIα LLPS remain unclear. Here, we reveal that two separate dimerization interfaces combined with the cAMP-induced release of the PKA catalytic subunit (PKA-C) from the pseudosubstrate inhibitory sequence are required to drive RIα condensate formation in cytosol, which is antagonized by docking to A-kinase anchoring proteins. Strikingly, we find that the RIα pseudosubstrate region is critically involved in the formation of a non-canonical R:C complex, which serves to maintain low basal PKA activity in the cytosol by enabling the recruitment of active PKA-C to RIα condensates. Our results suggest that RIα LLPS not only facilitates cAMP compartmentation but also spatially restrains active PKA-C, thus highlighting the functional versatility of biomolecular condensates in driving signaling specificity.

## INTRODUCTION

Just a handful of signaling pathways are tasked with coordinating distinct functional responses to the array of external cues encountered by cells. Among these, 3’,5’-cyclic adenosine monophosphate (cAMP) and its main effector cAMP-dependent protein kinase (PKA) transduce signals from hundreds of G-protein-coupled-receptors (GPCRs) to regulate cell metabolism, growth, and survival, as well as neuronal, cardiac, and pancreatic function^1–6^. This broad functional repertoire demands precise control of cAMP/PKA signaling specificity, which cells accomplish using numerous mechanisms. For example, the PKA holoenzyme comprises a pair of catalytic (C) subunits bound to one of four non-redundant, dimeric regulatory (R) subunits (RIα, RIβ, RIIα, RIIβ) expressed in different tissues^7,8^, while A-kinase anchoring proteins (AKAPs) recruit the PKA holoenzyme to discrete signaling compartments throughout the cell^9,10^. The heart of the PKA holoenzyme is the R:C complex, through which cAMP allosterically controls PKA catalytic activity^11,12^. In the canonical view of PKA activation, cooperative binding of two cAMP molecules to a pair of cyclic nucleotide-binding (CNB) domains initiates an allosteric switch that is transmitted through a flexible linker docked within the C subunit catalytic site and culminates in the release and dissociation of PKA-C^13^. Recent work alternatively suggests that PKA-C is unleashed but remains tethered to RIα and RII subunits^14,15^, though others failed to find direct supporting evidence for RI or RII^16^. It is therefore unclear whether or how such a noncanonical R:C complex can form and how it may impact PKA signaling.

Liquid-liquid phase separation (LLPS) is increasingly viewed as a critical organizer of the cell’s biochemical activity architecture^17–20^. We recently reported that the ubiquitously expressed RIα subunit of PKA undergoes LLPS to form biomolecular condensates in cells^21^. RIα condensates form in response to dynamic changes in cAMP levels and robustly sequester high concentrations of cAMP, both *in vitro* and in cells. This behavior is consistent with an emerging model in which physiological cAMP elevations are highly buffered, thereby slowing the effective diffusion of free cAMP and allowing cAMP-degrading phosphodiesterases (PDEs) to produce distinct cAMP signaling compartments^22^. Indeed, we found that the formation of RIα condensates in cells is required for PDE activity to successfully produce local cAMP-free nanodomains^21^, a key feature of cAMP compartmentation^23^. Disruption of RIα LLPS impaired PDE-mediated local cAMP degradation, leading to aberrant cAMP elevations and pathological signaling^21^. RIα LLPS thus appears to play a key role in driving cAMP compartmentation and signaling specificity. Nevertheless, little is known regarding the molecular details of RIα condensates, particularly the molecular mechanisms underlying RIα LLPS. Furthermore, we showed PKA-C is co-recruited to RIα condensates upon cAMP elevation, a condition that should in principle lead to dissociation of the PKA holoenzyme. The molecular mechanisms and functional role of this co-recruitment are not clear.

Here, we introduced mutations into RIα to target discrete functional units within the R:C complex and more precisely dissect the molecular determinants of RIα LLPS. We show that two distinct interfaces within the RIα dimer are required for LLPS, while both CNB domains mediate cAMP-induced condensate formation. Sequence analysis combined with mutations further shows that the amino acid composition of the flexible linker within RIα is important for LLPS. In addition, we provide evidence of a noncanonical R:C complex which allows active PKA-C to remain physically associated with RIα within condensates, leading to the sequestration of PKA-C and reduction of basal PKA activities in the cytosol. Our results shed new light on the mechanism and functional roles of RIα LLPS and the behavior of the R:C molecular machine.

## RESULTS

### Dimerization promotes and AKAP anchoring disrupts RIα condensate formation in cytosol

We previously found that deleting the N-terminal docking and dimerization (D/D) domain by truncating residues 12-61 (Fig. 1A) completely abolishes RIα condensate formation in cells^21^. However, as the D/D domain not only mediates R subunit dimerization but also provides the docking interface for interactions with AKAPs^9,24^, we sought to more carefully examine whether and how these two distinct functions contribute to RIα LLPS. The RI D/D domain comprises a set of α-helices that form an antiparallel dimer interface between opposing subunits^25,26^. The conserved residue L50 lies near the heart of the D/D domain and forms part of the hydrophobic core of the dimer interface (Fig. 1B). A single Leu-to-Arg substitution at this position (L50R), originally identified in RIβ as the pathological driver of a hereditary neurodegenerative disorder, is speculated to disrupt both RI dimer formation (Fig. 1B, Suppl. Table 1) and AKAP anchoring^27^. We therefore hypothesized that introducing the L50R mutation into RIα would allow us to more precisely assess the role of the D/D domain in promoting RIα LLPS.

**Figure 1.**
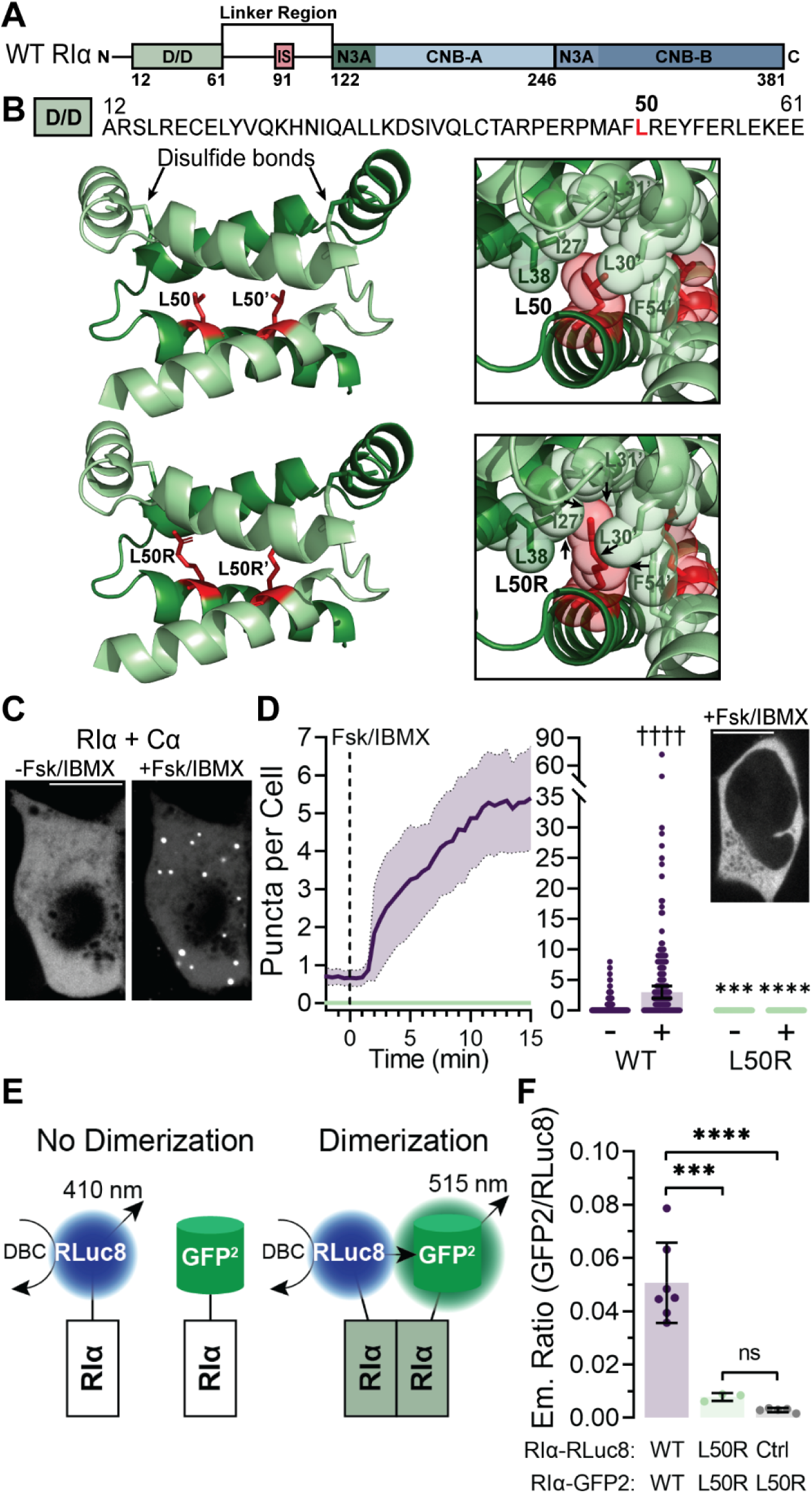
RIα D/D domain promotes LLPS via dimerization. **(A)** PKA RIα domain structure, including the Dimerization/Docking (D/D) domain, intrinsically disordered linker region, which includes the Inhibitory Sequence (IS), and the cyclic nucleotide-binding domains, CNB-A and CNB-B. **(B)** Amino acid sequence **(top)** and structural models (PBD: 3IM4 and L50R model) **(bottom)** of the RIα D/D domain. Highlighted: two disulfide bonds and residues involved in D/D dimerization, including L50 or L50R (red) and clashes (arrows). **(C)** Representative maximum intensity projections from confocal z-stacks showing GFP fluorescence from HEK293T cells co-expressing RIα-GFP2 plus Cα-mCherry before (−) and after (+) Fsk (50 µM) and IBMX (100 µM) stimulation. **(D)** Average time-course (left) and summary (right) of puncta number per cell before (−) and after (+) Fsk/IBMX stimulation in HEK293T cells expressing Cα-mCherry plus either wild-type (WT) RIα-GFP2 (WT; n = 157 cells from 4 experiments) or RIα^L50R^-GFP2 (L50R; n = 92 cells from 2 experiments). Inset shows a representative confocal fluorescence image of an RIα^L50R^-GFP2-expressing cell. Time-courses indicate the mean (solid line) and 95% confidence intervals (CI, shading). Error bars in summary quantification indicate median ± 95% CI. ††††P < 1 × 10^−15^ (WT+ vs WT-), paired Wilcoxon rank sum test; ***P = 0.000221 (L50R- vs WT-) and ****P < 1 × 10^−15^ (L50R+ vs WT+), unpaired Komogorov-Smirnov test. **(E)** BRET2 assay for monitoring RIα dimerization. **(F)** Maximum GFP2/RLuc8 emission ratio in HEK293T cells expressing RLuc8 and GFP2 fused to RIα WT or L50R: WT-RLuc8 with WT-GFP2 (WT/WT; n = 7 experiments), L50R-RLuc8 with L50R-GFP2 (L50R/L50R; n = 3 experiments), and RLuc8 alone (Ctrl) with L50R-GFP2 (Ctrl/L50R; n = 5 experiments). ***P = 0.000227 and ****P = 1.67 × 10^−5^ vs WT/WT; ordinary one-way ANOVA followed by Tukey’s multiple comparisons test. Error bars indicate mean ± SD. All scale bars, 10 µm.

We first investigated whether the L50R mutation affects RIα LLPS by comparing the ability of wild-type (WT) or L50R-mutant RIα to form droplets *in vitro*. Consistent with our previous study^21^, purified RIα formed liquid droplets *in vitro*, which was facilitated by cAMP (Suppl. Fig. 1A-B), likely due to increased disorder of the RIα linker region. Importantly, purified PKA-Cα inhibited RIα liquid droplet formation and cAMP counteracted the inhibitory effects of PKA-Cα on RIα phase separation, allowing liquid droplet formation at lower RIα concentrations (Suppl. Figure 1C), similar to our previous findings^21^. However, we were unable to purify sufficient quantities of recombinant RIα^L50R^ for these assays and therefore shifted to examining condensate formation with fluorescently tagged RIα and RIα^L50R^, which were both well expressed in HEK293T cells (Suppl. Fig. 1D).

As we observed previously^21^, HEK293T cells co-expressing GFP2-tagged WT RIα (RIα-GFP2) along with mCherry-tagged PKA-Cα (Cα-mCherry) exhibited no basal RIα condensate formation, with a median of 0 puncta per cell (95% confidence interval [CI]: 0-0; n = 157 cells) (mean ± SD; n = 54 cells), which increased to 3 puncta per cell (95% CI: 2-4; n = 157 cells; P < 1×10-15) with a partition of 6.26% ± 5.39% (n = 54 cells), upon treatment with the adenylyl cyclase activator forskolin (Fsk) and the PDE inhibitor 3-isobutyl-1-methylxanthine (IBMX) to induce cAMP accumulation (Fig. 1C-D, Suppl. Fig. 1E). In contrast to WT RIα, cells expressing RIα^L50R^-GFP2 plus Cα-mCherry exhibited no puncta formation either before or after Fsk/IBMX stimulation (P = 0.000221 (L50R- vs WT-) and P < 1×10^−15^ (L50R+ vs WT+); Fig. 1D), indicating that the single L50R point mutation is sufficient to recapitulate the LLPS-disrupting effect of the RIα D/D domain truncation.

Next, we investigated the molecular impact of the L50R mutation on PKA-R dimerization and R:C complex dissociation using bioluminescence resonance energy transfer (BRET)-based assays in living cells^28^. HEK293T cells co-transfected with GFP2-Cα plus *Renilla* luciferase (RLuc8)-tagged WT or L50R-mutant RIα exhibited similar decreases in the GFP-to-RLuc8 emission ratio upon Fsk stimulation (Suppl. Fig. 1F), indicating that the lack of stimulated puncta formation was not caused by loss of cAMP-responsiveness (Suppl. Fig. 1G-H). We then co-transfected HEK293T cells with pairs of plasmids encoding WT RIα or RIα^L50R^ tagged with GFP2 or RLuc8, such that the formation of GFP2-and-RLuc8-containing RIα heterodimers will lead to increased energy transfer from RLuc8 to GFP2, allowing the BRET ratio to serve as a proxy for RIα dimer formation (Fig. 1E). Compared with cells expressing WT RIα, which exhibited an emission ratio (R) of 0.0507 ± 0.0151 (mean ± SD; n = 7 experiments), cells expressing RIα^L50R^ constructs (L50R-RLuc8 + L50R-GFP2) exhibited a substantially lower emission ratio (R = 0.00786 ± 0.00150, n = 3 experiments; P = 0.000227) (Fig. 1F). Similarly low ratios were recorded from negative controls cells expressing L50R-GFP2 plus untagged RLuc8 (R = 0.00293 ± 0.000775, n = 5 experiments; ns, not significant), suggesting that the low emission ratios measured in cells expressing L50R-RLuc8 + L50R-GFP2 were due to the loss of RIα dimer formation.

As the L50R mutation is also expected to impact AKAP binding, we further examined the impact of AKAP docking on RIα LLPS by monitoring the effect of AKAP co-expression on RIα subcellular localization and puncta formation. When we co-expressed mRuby2-tagged WT RIα in HEK293T cells along with EGFP-tagged smAKAP, an RI-specific AKAP known to localize to the plasma membrane (PM)^29^, we observed strong PM colocalization of RIα-mRuby2 and smAKAP-EGFP fluorescence, consistent with smAKAP-mediated membrane recruitment (Fig. 2A). Importantly, whereas WT RIα-mRuby2 successfully formed puncta when expressed alone, no RIα-mRuby2 puncta were detected when smAKAP-EGFP was co-expressed (Suppl. Fig. 2A-B). In contrast, RIα^L50R^-mRuby2 fluorescence was diffusely distributed and showed no colocalization with smAKAP-EGFP (n = 10 cells from 3 independent experiments) (Fig. 2A), confirming that the L50R mutation disrupts AKAP anchoring of RIα. RIα^L50R^-mRuby2 also failed to form puncta in cells co-expressing smAKAP-EGFP, consistent with our earlier observations (Fig. 1D, Suppl. Fig. 2B). Interestingly, RIα-mRuby2 puncta formation was observed upon co-expression with smAKAP in which L65 and L70 were mutated to Pro (smAKAP^L2P2^-EGFP) (n = 10 cells from 3 independent experiments) (Fig. 2A, Suppl. Fig. 2B), which abolishes RI recruitment to AKAP (Suppl. Fig. 2C). These data suggest that loss of LLPS by RIα^L50R^ is likely not due to the loss of AKAP binding, as AKAP anchoring appears to inhibit RIα condensate formation in cytosol. This inhibitory effect could be achieved by either 1) AKAP binding to the RIα D/D domain or 2) sequestration of RIα from the diffusible, cytosolic pool.

**Figure 2.**
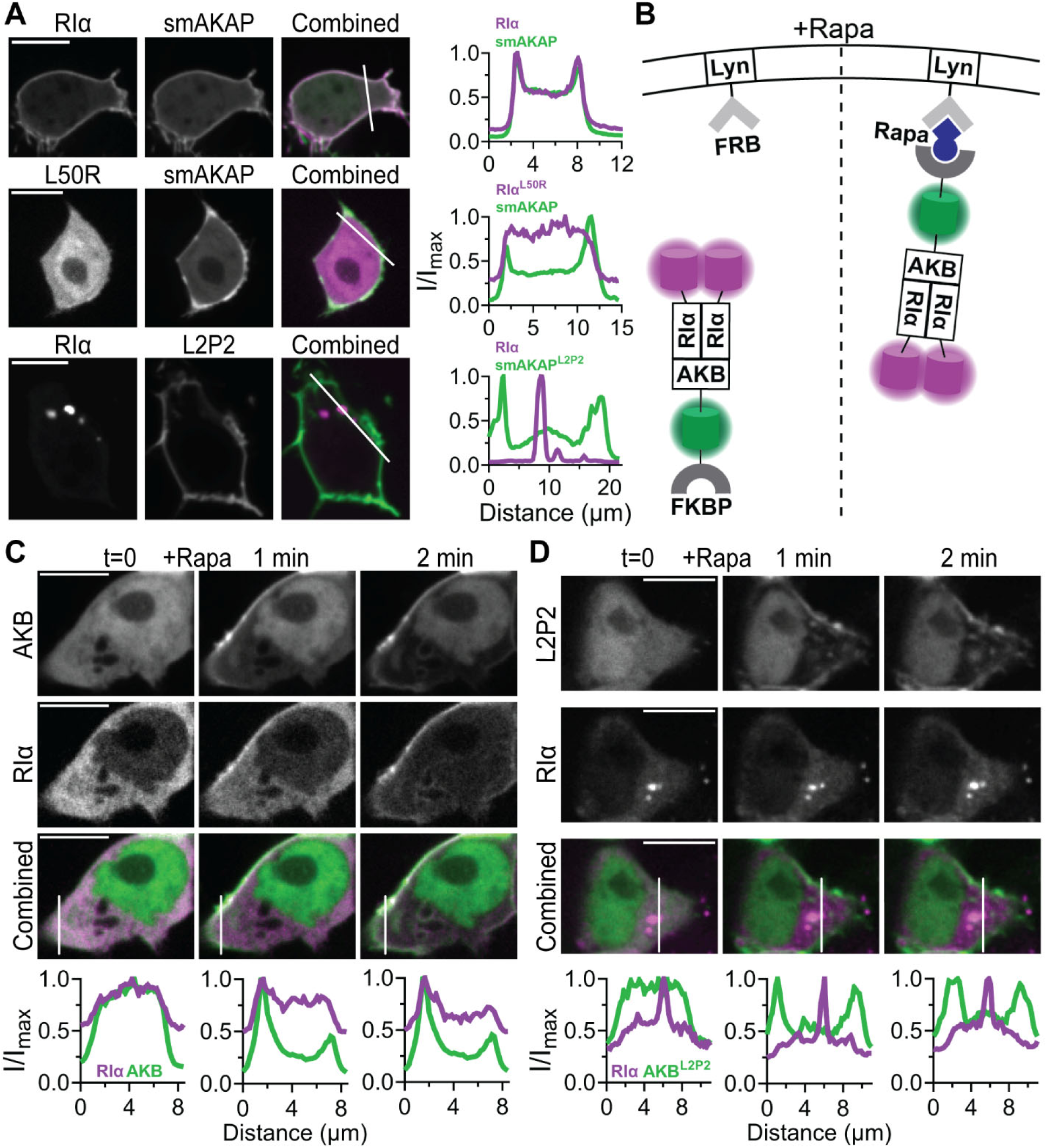
AKAP docking disrupts RIα condensate formation in cytosol. **(A)** Representative confocal fluorescence images and line-intensity profiles of the indicated regions of HEK293T cells co-expressing EGFP-tagged wild-type (WT) smAKAP or smAKAP^L2P2^ (L2P2) plus mRuby2-tagged WT or L50R-mutant RIα (n = 10 cells from 4 independent experiments). **(B)** Recruitment of RIα-bound A-kinase-binding (AKB) domain to the plasma membrane using the rapamycin (Rapa)-induced dimerization of FKBP and FRB. **(C-D)** Representative confocal fluorescence images and line-intensity profiles of the indicated regions of HEK293T cells co-expressing **(C)** WT or **(D)** L2P2-mutant AKB-mVenus-FKBP plus RIα-mRuby2 and Lyn-FRB before (t = 0) as well as 1 and 2 min after addition of 1 µM rapamycin (n = 10 cells from 4 independent experiments). All scale bars, 10 µm.

To investigate the former possibility, we designed an approach based on the chemically inducible dimerization of FKBP and FRB^30,31^. We fused the isolated A-kinase-binding (AKB) domain of smAKAP to mVenus and FKBP (AKB-mVenus-FKBP), which we co-expressed with PM-targeted FRB (Lyn-FRB) and RIα-mRuby2, allowing us to acutely induce the PM translocation of AKB, and by extension RIα, through the addition of rapamycin (Fig. 2B). Notably, we found no RIα-mRuby2 puncta in cells co-expressing AKB-mVenus-FKBP compared with control cells co-expressing AKB^L2P2^-mVenus-FKBP (0 [95% CI: 0-0] vs. 4 [95% CI: 2-3] puncta per cell, P < 1×10^−15^; n = 59 and 60 cells, respectively), suggesting that D/D-domain-mediated AKAP docking to the RIα dimer is sufficient to abolish RIα condensate formation in cytosol (Suppl. Fig. 2B). Rapamycin addition further resulted in re-localization of the AKB-mVenus-FKBP and RIα-mRuby2 fluorescence signals to the PM (Fig. 2C, Suppl. Fig. 2D), confirming the AKB-RIα interaction, whereas RIα-mRuby2 fluorescence did not re-localize in AKB^L2P2^-mVenus-FKBP cells (Fig. 2D, Suppl. Fig. 2D). Altogether, our results demonstrate that the D/D domain drives RIα LLPS exclusively through mediating RIα dimerization and further suggest that RIα LLPS and AKAP anchoring are distinct forms of spatial compartmentation within the cAMP/PKA pathway.

### RIα LLPS involves a secondary dimer interface

The N3A motif, a short helical motif comprising an N-helix, 3_10_-loop, and A-helix, is a conserved structural feature located at the N-terminus of CNB domains in various cAMP-regulated proteins^32^. In addition to the established role of this motif in the cAMP-triggered allosteric switch^32^, structural analysis of RIα revealed that the N3A motif of CNB-A forms a secondary dimer interface that is unique to RI isoforms^33–35^. This N3A dimer interface consists of a tightly packed helical bundle, where key residues such as Y122 (corresponding to Y120 in bovine RIα) form critical π-stacking and hydrogen-bond interactions between opposing protomers. Ala substitution of Y122 has previously been shown to disrupt the typically compact conformation of the RIα homodimer (Fig. 3A)^33^. Given the importance of multivalent interactions in driving biomolecular phase transitions^18,36,37^, we introduced the Y122A mutation into human RIα to investigate whether interactions mediated by the CNB-A N3A motif interface also contribute to RIα LLPS.

**Figure 3.**
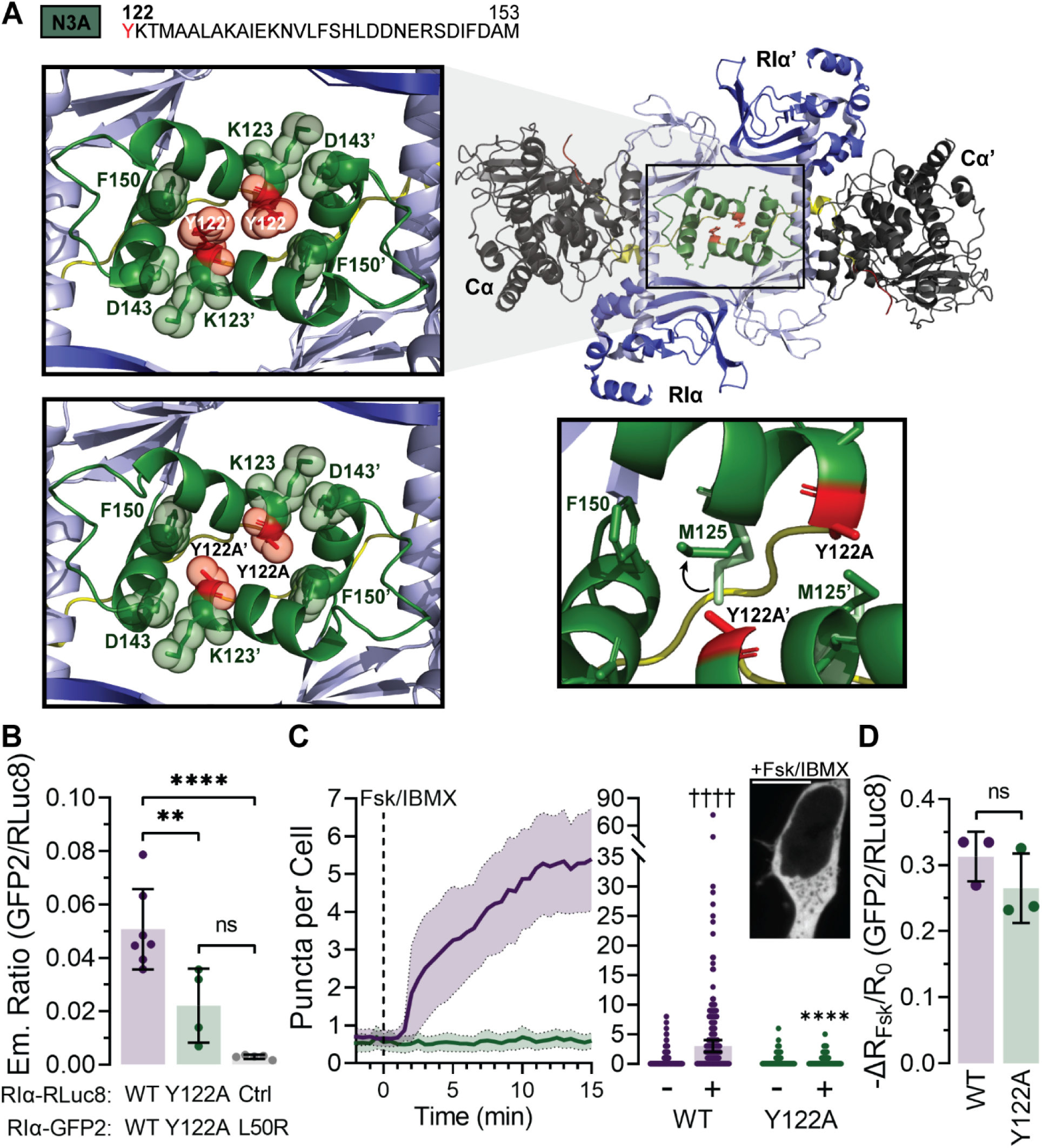
Secondary dimerization via the CNB-A N3A motif promotes RIα LLPS. **(A)** Amino acid sequence **(top)** and crystal structures (PBD: 6BYS and Y122A model) **(bottom)** of the RIα CNB-A N3A motif (green) dimerization at the C-terminus of linker region (yellow). RIα CNB-A is light blue and CNB-B is dark blue. Highlighted: residues involved in CNB-A N3A motif dimerization, including Y122 and Y122A (red). **(B)** Maximum GFP2/RLuc8 emission ratio in HEK293T cells expressing RLuc8 and GFP2 fused to RIα WT or Y122A: WT-RLuc8 with WT-GFP2 (WT/WT; data reproduced from Fig. 1), Y122A-RLuc8 with Y122A-GFP2 (Y122A/Y122A; n = 4 experiments), and RLuc8 alone (Ctrl) with L50R-GFP2 (Crtl/L50R; data reproduced from Fig. 1). **P = 0.00656 and ****P = 4.24 × 10^−5^ vs WT/WT; ordinary one-way ANOVA followed by Tukey’s multiple-comparisons test. Error bars indicate mean ± SD. **(C**) Average time-course (left) and summary (right) of puncta number per cell before (−) and after (+) Fsk/IBMX stimulation in HEK293T cells co-expressing Cα-mCherry plus either RIα-GFP2 (WT, reproduced from Fig. 1) or RIα^Y122A^-GFP2 (Y122A, n = 95 cells from 2 experiments). Inset shows a representative confocal fluorescence image of an RIα^Y122A^-GFP2-expressing cell. ††††P < 1 × 10^−15^ (WT+ vs WT-), paired Wilcoxon rank sum test; ****P = 2.48 × 10^−12^ (Y122A+ vs WT+), unpaired Komogorov-Smirnov test. Time-courses indicate the mean (sold line) ± 95% CI (shading). Error bars in summary quantification indicate median ± 95% CI. Scale bar, 10 µm. **(D)** Fsk-stimulated change in the GFP2/RLuc8 emission ratio in HEK293T cells co-expressing GFP2-Cα plus RLuc8 fused to either WT RIα or RIα^Y122A^. n = 3 experiments each. ns, not significant; unpaired, two-tailed Student’s t-test. Error bars indicate mean ± SD.

We probed the effect of the Y122A mutation on RIα using the previously described BRET dimerization assay and found that HEK293T cells co-expressing Y122A-GFP2 and Y122A-RLuc8 exhibited a lower emission ratio (R = 0.022 ± 0.014, n = 4 experiments) compared with cells co-expressing GFP2- and RLuc8-tagged WT RIα (R = 0.0507 ± 0.0151, n = 7 experiments, P = 0.00656 vs Y122A, Fig. 3B). Although structural models show that the Ala substitution of Y122 does not introduce any steric clashes (Fig. 3A), unlike the L50R mutation (Suppl. Table 1), our results confirm that disrupting the CNB-A N3A motif interferes with RIα dimerization, in agreement with previous biophysical characterization^33^.

Consistent with this effect, HEK293T cells co-expressing RIα^Y122A^-GFP2 plus Cα-mCherry showed reduced puncta formation compared with WT RIα (Fig. 3C). In particular, we observed a complete loss of cAMP-stimulated puncta formation upon treatment with Fsk/IBMX (0 [95% CI: 0-0] vs. 3 [95% CI: 2-4] puncta per cell for Y122A vs. WT, P = 2.48 ×10^−12^; n = 95 and 157 cells, respectively). The lack of cAMP-induced puncta formation by RIα^Y122A^ was not due to an impaired cAMP response, as HEK293T cells expressing GFP2-Cα plus either WT RIα- or RIα^Y122A^-RLuc8 showed similar Fsk-stimulated emission ratio changes (-ΔR_Fsk_/R_0_ = 31.3 ± 3.76% and 26.5 ± 05.25% for WT and Y122A, respectively, n = 3 experiments; ns, not significant) (Fig. 3D, Suppl. Fig. 3). Our results thus implicate multiple interaction sites in driving RIα LLPS.

### Both RIα CNB domains are required for cAMP induced LLPS

We have shown that cAMP plays a critical role in regulating the formation of RIα biomolecular condensates^21^. Binding of cAMP to RIα is mediated by a pair of cooperative CNB domains that exhibit distinct affinities for cAMP^11,38^. The CNB-A domain shows a 50-fold higher EC_50_ for cAMP compared with the CNB-B domain, and cAMP is thought to first bind to CNB-B, resulting in increased affinity of CNB-A for cAMP^39,40^. While our prior truncation study suggested that RIα LLPS requires the CNB domains^21^, we sought to test the individual contribution of cAMP binding to each CNB domain to RIα LLPS.

Residues E202 and R211 of human RIα (corresponding to E200 and R209, respectively, in bovine RIα) form key hydrogen bonds that coordinate cAMP binding within the CNB-A domain (Fig. 4A)^41,42^. Similarly, residues E326 and R335 (corresponding to E324 and R333, respectively, in bovine RIα), in conjunction with four other residues, form hydrogen bonds with cAMP within the CNB-B domain (Fig. 4B)^39,43^. To precisely test the role of each CNB domain in cAMP-induced RIα LLPS, we mutated either E202 or E326 to Ala. BRET-based assays revealed that neither RIα^E202A^- nor RIα^E326A^-RLuc8 yielded a Fsk-induced change in the emission ratio when co-expressed with GFP2-Cα in HEK293T cells, in contrast to WT RIα-RLuc8, confirming that each mutation was sufficient to render RIα unresponsive to cAMP (Fig. 4C, Suppl. Fig. 4A). Consistent with these results, we found that both RIα^E202A^- and RIα^E326A^-GFP2, although capable of forming puncta when expressed alone (Suppl. Fig. 4B-C), showed no cAMP-induced puncta formation in HEK293T cells co-expressing Cα-mCherry and treated with Fsk/IBMX (RIα^E202A^-GFP2: 0 [95% CI: 0-0] puncta per cell, n = 118 cells, P < 1×10^−15^ vs. WT; RIα^E326A^-GFP2: 0 [95% CI: 0-0] puncta per cell, n = 89 cells, P < 1×10^−15^) (Fig. 4D). These results suggest that cAMP regulation of RIα phase separation requires binding to both CNB domains.

**Figure 4.**
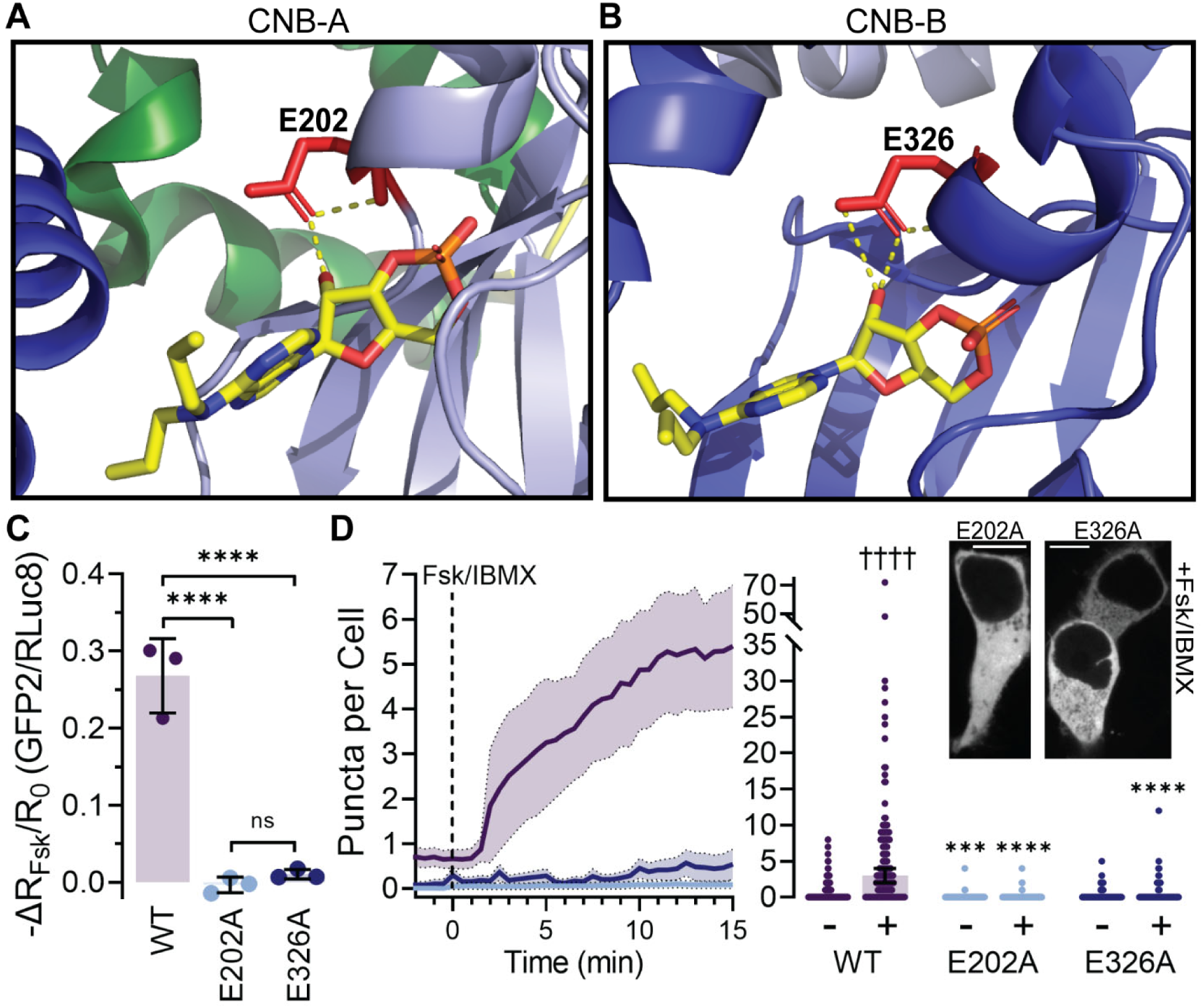
Both CNB domains are required for cAMP-induced RIα LLPS. Crystal structures (PBD: 4JV4) of cAMP bound to the **(A)** CNB-A and **(B)** CNB-B of RIα, highlighting hydrogen bonds formed with residues **(A)** E202 and **(B)** E326. **(C)** Fsk-stimulated change in the GFP2/RLuc8 emission ratio in HEK293T cells co-expressing GFP2-Cα plus RLuc8 fused to WT RIα, RIα^E202A^, or RIα^E326A^. n = 3 experiments each. ****P = 5.97 × 10^−5^ (E202A vs WT) and 8.08 × 10^−5^ (E326A vs WT); ordinary one-way ANOVA followed by Tukey’s multiple-comparisons test. Error bars indicate mean ± SD. **(D**) Average time-course (left) and summary (right) of puncta number per cell before (−) and after (+) Fsk/IBMX stimulation in HEK293T cells co-expressing Cα-mCherry plus WT RIα (WT; reproduced from Fig. 1), RIα^E202A^ (E202A; n = 118 cells from 2 experiments), and RIα^E326A^ (E326A; n = 89 cells from 2 experiments). Insets show representative confocal fluorescence images of HEK293T cells expressing RIα^E202^- or RIα^E326A^-GFP2. Time-courses indicate the mean (solid lines) ± 95% CI (shading). Error bars in summary quantification indicate median ± 95% CI. ††††P < 1 × 10^−15^ (WT+ vs WT-); paired Wilcoxon rank sum test and ***P = 0.000317 (E202A- vs WT-), ****P < 1 × 10^−15^ (E202A+ vs WT+), and ****P < 1 × 10^−15^ (E326A+ vs WT+); unpaired Komogorov-Smirnov test. Scale bars, 10 µm.

### RIα LLPS is a function of IS binding state

Our *in vitro* studies indicate that the presence of PKA-C inhibits RIα LLPS, which can be reversed by cAMP (Suppl. Fig. 1C)^21^. The inability of CNB domain mutants to dissociate from Cα or form cAMP-induced puncta (Fig. 4C-D) further supports a model in which the unleashing of PKA-C is required for RIα to phase separate. We previously found that removing the RIα linker region via deletion of residues 62-113 (Fig. 1A) abrogated RIα LLPS, highlighting the importance of this region^21^.

The linker region is not resolvable in crystal structures of the free RIα dimer^33^, suggesting that this region is highly disordered. Indeed, sequence analysis predicts a high degree of disorder, especially within N-terminal residues 62-90, which are highly (76.7%) enriched in disorder-promoting residues (P, E, K, S and Q) (Suppl. Fig. 5A-C)^44–46^, consistent with the key role of intrinsic disorder in promoting phase separation^18,36,37^. Furthermore, the sticker and spacer model of LLPS holds that charged residues (K, R, D, and E, coral in Figure 5A) help drive phase-separation behavior by conferring both sticker- and spacer-like characteristics of interaction affinity and solvation volume to intrinsically disordered regions^36,47–49^. Net charge per residue (NCPR) has been adopted as a metric to illustrate the effect of charged residues on phase separation^50^ and mutating a large number of charged residues to Ala is calculated to shift the NCPR of the RIα linker region from 0.05 to 0.083 (Suppl. Fig. 5D). This trend is predicted to disrupt LLPS, and indeed, the charge-to-Ala (Ch2A) mutant abolished RIα puncta formation in cells in both the presence and absence of PKA-C (0 [95% CI: 0-0] puncta per cell without PKA-C, n = 512 cells, P < 1×10^−15^ vs. WT; 0 [95% CI: 0-1] puncta per cell to 1 [95% CI: 0-2] puncta per cell with PKA-C, n = 60 cells, P = 9.08×10^−10^ (Ch2A+ vs WT+)) (Suppl. Fig. 5E).

**Figure 5.**
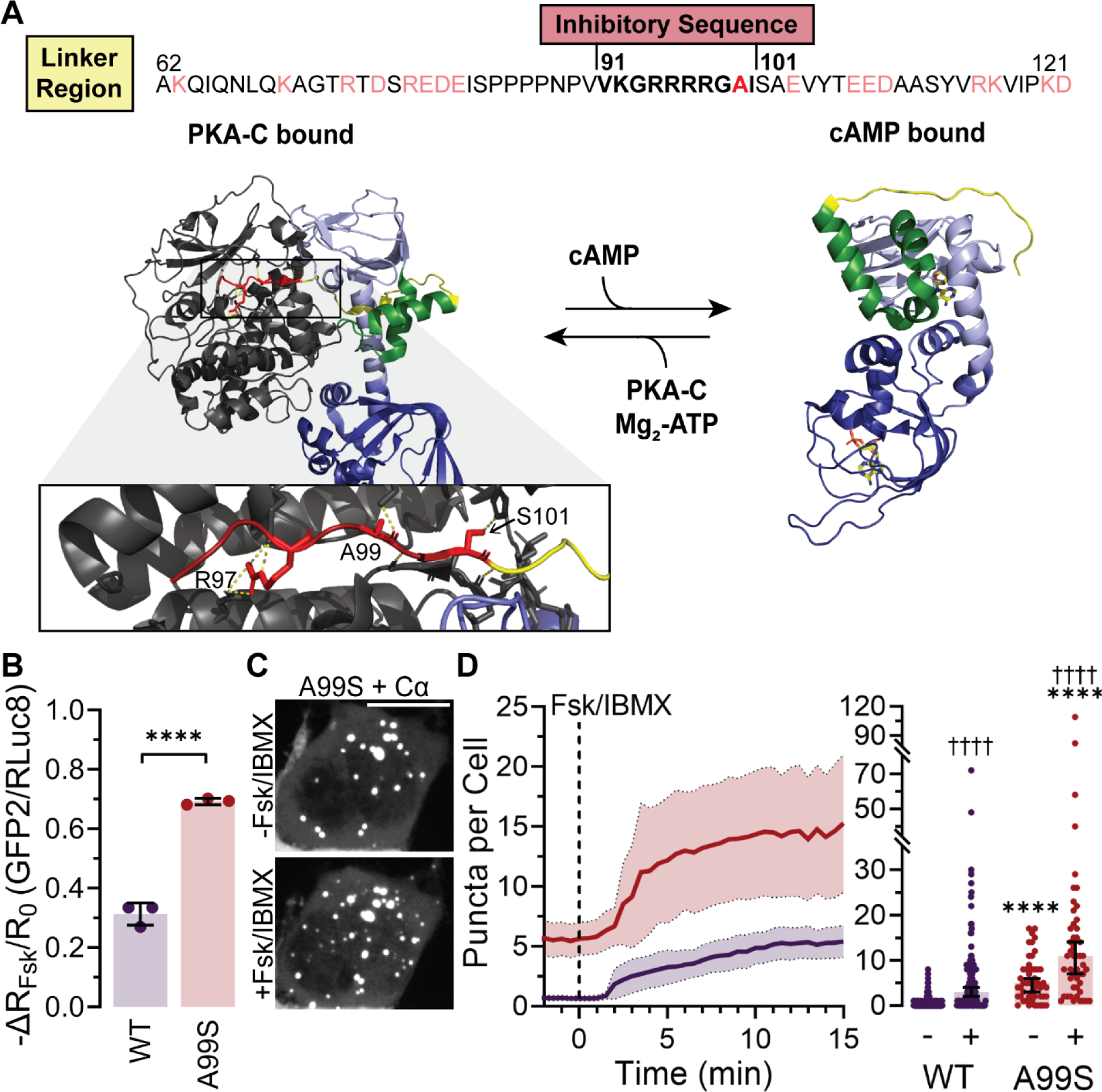
RIα LLPS is a function of IS binding state. **(A)** Amino acid sequence of the RIα linker region, including the linker region charged residues (coral), Inhibitory Sequence (IS) (bold), and the A99 “P”-site (red) **(top)** and crystal structure of PKA-C-(left; PBD: 6NO7) or cAMP-bound (right; PBD: 4MX3) RIα, highlighting RIα residues that interact with Cα residues **(bottom)**. In the structures, the PKA-C is grey and the RIα linker is yellow, IS is red, CNB-A N3A motif is green, CNB-A is light blue and CNB-B is dark blue. **(B)** Fsk-stimulated change in the GFP2/RLuc8 emission ratio in HEK293T cells co-expressing GFP2-Cα plus RLuc8 fused to either WT RIα (WT; data reproduced from Fig. 3) or RIα^A99S^ (A99S; n = 3 experiments). ****P = 7.33 × 10^−5^ vs WT; unpaired, two-tailed Student’s t-test. Error bars indicate mean ± SD. **(C)** Representative maximum intensity projections from confocal z-stacks of HEK293T cells co-expressing Cα-mCherry and RIα^A99S^-GFP2 in HEK293T cells. Scale bar, 10 µm. **(D)** Average time-course (left) and summary (right) of puncta number per cell before (−) and after (+) Fsk/IBMX stimulation in HEK293T cells co-expressing Cα-mCherry plus either RIα-GFP2 (WT; data reproduced from Fig. 1) or RIα^A99S^-GFP2 (A99S; n = 49 cells from 2 experiments). Time-courses indicate the mean (solid lines) ± 95% CI (shading). Error bars in summary quantification indicate median ± 95% CI. ††††P < 1 × 10^−15^ (WT+ vs WT-) and ††††P = 1.86 × 10^−8^ (A99S+ vs A99S-); paired Wilcoxon rank sum test and ****P = 4.71 × 10^−13^ (A99S- vs WT-) and ****P = 7.83 × 10^−6^ (A99S+ vs WT+); unpaired Komogorov-Smirnov test.

When PKA-C is bound to RIα, the inhibitory sequence (IS) within the linker region is buried near the PKA-C active site cleft (Fig. 5A, left)^13^. cAMP binding to both CNB domains induces conformational changes that allosterically signal the unleashing of PKA-C, increasing exposure of the linker (Fig. 5A, right)^13^. Our results are consistent with a model in which the LLPS-promoting characteristics of the RIα linker are suppressed by PKA-C binding and cAMP-induced exposure and subsequent disordering of the linker drives cAMP-induced RIα LLPS. We therefore hypothesized that IS binding state directly controls RIα phase separation. In RIα, the IS comprises a pseudosubstrate that inhibits PKA-C catalytic activity when bound to PKA-C within the holoenzyme^13^. Mutating the putative phosphorylation site (P^0^) from Ala to Ser (A99S) has been shown to trigger RIα autophosphorylation^51,52^ and enhance cAMP-induced dissociation of RIα and PKA-C, suggesting a weakened R:C complex (Fig. 5A)^53^. Consistent with these reports, we observed in BRET assays that HEK293T cells co-expressing GFP2-Cα plus RIα^A99S^-RLuc8 exhibited a much larger Fsk-induced emission ratio change (-ΔR_Fsk_/R_0_) of 69.2 ± 1.08% (n = 3 experiments), compared with 31.3 ± 3.76% (n = 3 experiments) for WT RIα-RLuc8 (P = 7.33×10^−5^) (Fig. 5B, Suppl. Fig. 5F). Similarly, the phosphomimetic A99E RIα mutant also showed a large Fsk-induced emission ratio change of 69.7 ± 11.4% (Suppl. Fig. 5F-G)^53^. These data suggest that phosphorylation of the P-site in A99S promotes the release of Cα by RIα and thus exposure of the disordered linker.

When we examined the impact of this mutation on RIα LLPS, we found that co-expressing RIα^A99S^-GFP2 along with Cα-mCherry in HEK293T cells resulted in substantially higher levels of puncta formation, both basally (Fig. 5C-D; 5 [95% CI: 3-6] puncta per cell, n = 49 cells; P = 4.71×10^−13^ vs. WT) and following cAMP elevation using Fsk/IBMX (11 [95% CI:7-14] puncta per cell, n = 49 cells; P = 7.83×10^−6^ vs. WT), indicating that weakening the R:C interaction enhances RIα LLPS. Comparable results were observed using the A99E RIα mutant (Suppl. Fig. 5H), suggesting that the increase in RIα puncta formation in both the basal and stimulated states may result from autophosphorylation of the IS, consistent with the previously observed autophosphorylation of an A99S mutant^51^. *In vitro*, purified A99S and Cα showed biphasic droplet formation (Suppl. Fig. 5I), with Cα exerting an inhibitory effect at low concentrations, similar to wildtype RIα (Suppl Figure 1C), but a positive effect at high concentrations, presumably caused by increased Cα-mediated phosphorylation of A99S. Consistent with this model, droplet formation by the A99E mutant did not show this biphasic behavior and was largely insensitive to Cα (Suppl. Fig. 5I). Thus, phosphorylation of the P-site or a phosphomimetic mutation (A99E) within the RIα IS prevents Cα from inhibiting RIα LLPS.

As RIα condensates have been shown to both form and disperse in conjunction with increasing and decreasing cAMP concentrations^21^, we also stimulated cells using the β-adrenergic receptor agonist isoproterenol (Iso) to induce transient cAMP elevations (Suppl. Fig. 5J). Under these conditions, we found that the formation of cAMP-induced RIα puncta in cells expressing Cα-mCherry + RIα^A99S^-GFP2 was sustained (Suppl. Fig. 5K), further indicating disruption of R:C interaction via the altered IS. Collectively, these data support our model that exposure of the disordered linker region via release of the IS by PKA-C drives cAMP-induced RIα LLPS.

### Non-canonical PKA Cα recruitment to RIα condensates

When co-expressed in HEK293T cells, RIα-GFP2 and Cα-mRuby2 can be seen to colocalize within puncta following Fsk/IBMX treatment, indicated by an increase in the Pearson’s correlation coefficient (Fig. 6A-B, Suppl. Fig. 6A-B). These results align with our previous findings that RIα biomolecular condensates recruit active PKA-C^21^. In contrast, co-expressing RIα^A99S^-GFP2 with Cα-mRuby2 markedly reduced Cα colocalization to RIα puncta after Fsk/IBMX stimulation (Fig. 6A-B, Suppl. Fig. 6A-B), suggesting that PKA-C recruitment to RIα condensates is dependent on its ability to bind the IS pseudosubstrate region. Indeed, we found that co-expressing RIα-GFP2 and Cα-mRuby2 with RIα (62-113)-mTagBFP, which encodes the RIα linker alone, disrupted PKA-C colocalization with RIα puncta in cells stimulated with Fsk/IBMX (Suppl. Fig. 6C). Thus, the isolated RIα linker region is sufficient to compete for PKA-C binding such that PKA-C is no longer retained in RIα puncta following cAMP elevation.

**Figure 6.**
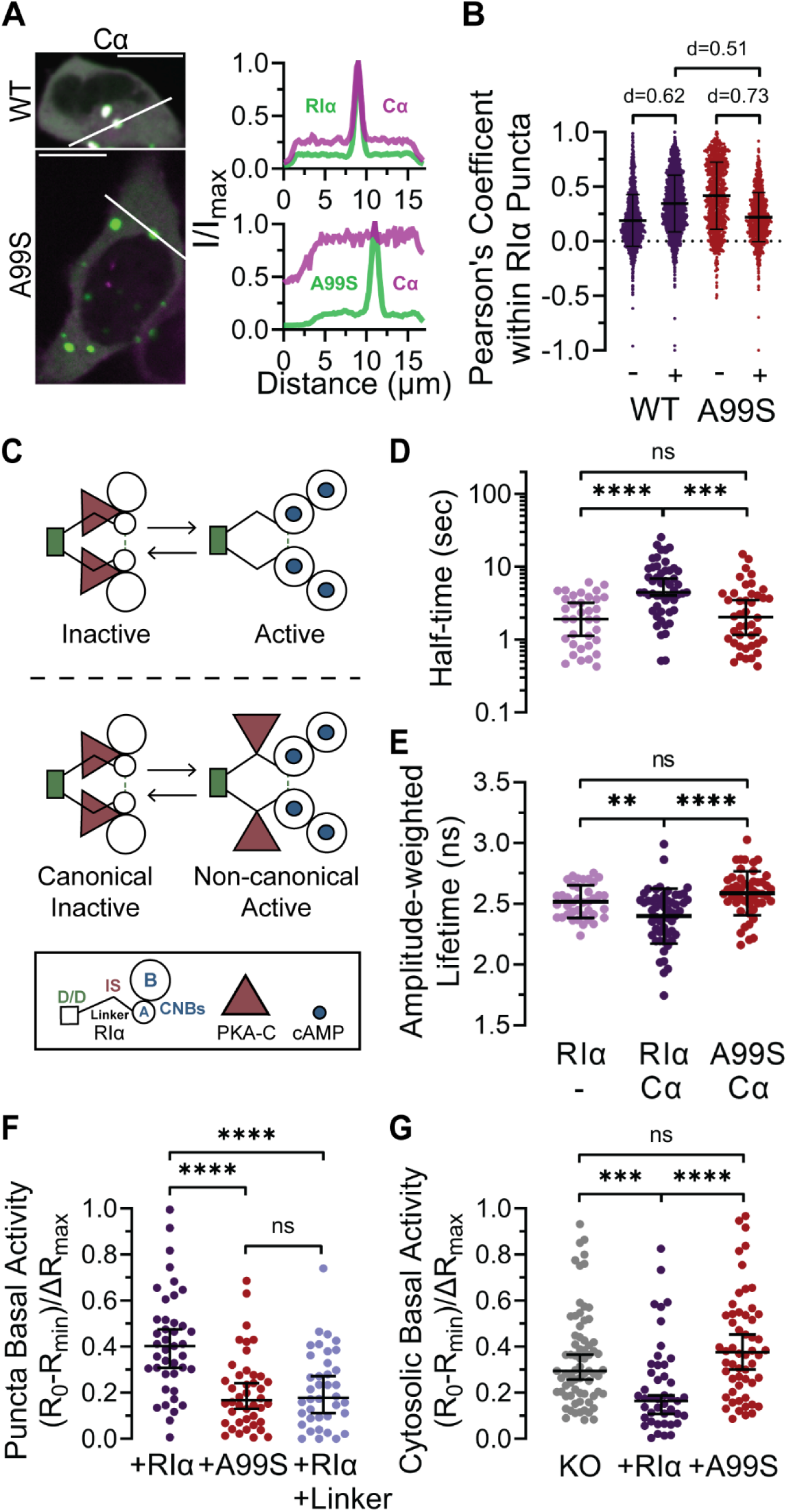
Non-canonical recruitment of PKA catalytic subunit to RIα condensates. **(A)** Representative confocal images and line-intensity profiles of the indicated regions of HEK293T cells co-expressing RIα-GFP2 plus Cα-mRuby2 (n = 10 from 4 independent experiments). Scale bars, 10 µm. **(B)** Pearson’s coefficient per RIα puncta in HEK293T cells co-expressing RIα-GFP2 (n = 2288 puncta before and n = 2502 puncta after stimulation) or RIα^A99S^-GFP2 (n = 1055 puncta before and n = 1253 after stimulation) with Cα-mRuby2. Effect size was calculated using Cohen’s d. Error bars indicate mean ± SD. **(C)** Schematic illustrating the classical model of dissociation of the canonical PKA holoenzyme (top) and the proposed model of the non-canonical R:C interaction (bottom). In the classical model, binding of cAMP (blue dots) to the CNB domains (open circles) causes complete dissociation of PKA-C (triangles), whereas our data support a non-canonical interaction in which active PKA-C remains tethered to RIα via the linker region. **(D)** Recovery kinetics (t_1/2_) of RIα puncta in cells expressing RIα-GFP2 alone (RIα/-; n = 35 puncta) or co-expressing Cα-mCherry plus either RIα-GFP2 (RIα/Cα; n = 51 puncta), or RIα^A99S^-GFP2 (A99S/Cα; n = 42 puncta) measured using FRAP. Error bars indicate median ± 95% CI. ****P = 6.15 × 10^−6^ (RIα vs RIα/Cα) and ***P = 0.000845 (RIα/Cα vs A99S/Cα); Brown-Forsythe and Welch ANOVA followed by Dunnett’s T3 multiple-comparisons test. **(E)** Average amplitude-weighted lifetimes measured from RIα puncta in HEK293T cells expressing RIα-GFP2 alone (RIα/-; n = 39 cells) or co-expressing Cα-mCherry plus either RIα-GFP2 (RIα/Cα; n = 54 cells) or RIα^A99S^-GFP2 (A99S/Cα; n = 50 cells). Error bars indicate mean ± SD. **P = 0.0066 (RIα/- vs RIα/Cα) and ****P = 2.91 × 10^−5^ (RIα/Cα vs A99S/Cα); Brown-Forsythe and Welch ANOVA followed by Dunnett’s T3 multiple-comparisons test. **(F)** Normalized FluoSTEP-AKAR R/G emission ratio when RIα-FP11 (RIα; n = 41 cells), RIα^A99S^-FP11 (A99S; n = 42 cells), or RIα-FP11 and RIα(62-113)-mTagBFP (RIα/Linker; n = 38 cells) are expressed in RIα-KO HEK293T cells treated with 10 µM H89, followed by washout and addition of Fsk/IBMX. ****P = 1.98 × 10^−5^ (RIα vs A99S) and ****P = 6.95 × 10^−5^ (RIα vs RIα/Linker); unpaired, two-tailed Student’s t-test. **(G)** Normalized AKAR4 Y/C emission ratio change in RIα-KO HEK293T cells treated with 10 µM H89, followed by washout and addition of Fsk/IBMX in the absence (KO; n = 69 cells) or presence of RIα-mRuby2 (RIα; n = 53 cells) or RIα^A99S^-mRuby2 (A99S; n = 87 cells) expression. Error bars indicate median ± 95% CI. ***P = 6.16 × 10^−4^ (KO vs RIα) and **** P = 1.23 × 10^−5^ (RIα vs A99S); Brown-Forsythe and Welch ANOVA followed by Dunnett’s T3 multiple-comparisons test.

Given that PKA-C association with RIα condensates was observed under high cAMP conditions and that high PKA activity was observed within the condensates^21^, we hypothesized that a non-canonical, catalytically active R:C complex (Fig. 6C) is formed within RIα condensates. In our model, this noncanonical complex is mediated by the pseudosubstrate IS, which is also critical for the canonical R:C complex. Yet the signaling capabilities of these two types of complexes are drastically different – whereas the canonical PKA holoenzyme is in an inhibited state, the noncanonical R:C complex minimizes the mutual “inhibition” between PKA-C and RIα so that PKA-C is active and RIα is able to form condensates. To investigate the possible existence of such a noncanonical R:C complex, we performed fluorescence recovery after photobleaching (FRAP) experiments to determine whether the ability of RIα to bind Cα via the linker region affects the exchange dynamics of RIα molecules. Quantification of puncta bleaching curves in cells co-transfected with RIα-GFP2 and Cα-mCherry revealed recovery kinetics (t_1/2_) of 6.66 ± 5.43 sec (n = 51 puncta), whereas cells co-expressing RIα^A99S^-GFP2 plus Cα-mCherry exhibited faster recovery kinetics of 3.20 ± 3.26 sec (n = 42 puncta; P = 0.000845) (Fig. 6D, Suppl. Fig 6D-E). Results from RIα^A99S^-containing puncta were comparable to those measured from puncta formed in cells expressing RIα alone (t_1/2_ = 2.42 ± 1.66 sec, n = 35 puncta; ns, not significant vs. RIα^A99S^-GFP2 + Cα-mCherry; P = 6.15×10^−6^ vs. WT RIα-GFP2 + Cα-mCherry), suggesting that the presence of Cα in the condensates slows down the exchange dynamics of WT RIα, possibly due to R:C interactions.

As a more direct measure of a putative non-canonical R:C complex within RIα condensates, we also performed fluorescence lifetime imaging microscopy-fluorescence resonance energy transfer (FLIM-FRET) measurements on RIα puncta formed in RIα-GFP2 and Cα-mCherry co-transfected cells following stimulation with Fsk/IBMX. Similar to the BRET assays described above, any interaction between RIα-GFP2 and Cα-mCherry within puncta is expected to increase FRET between the GFP2 donor and mCherry acceptor, thereby decreasing the donor fluorescence lifetime. Consistent with the FRAP data, we observed significantly longer donor fluorescence lifetimes within puncta from RIα^A99S^-GFP2 + Cα-mCherry cells (τ = 2.59 ± 0.182 ns, n = 50 cells) compared with puncta in WT RIα-GFP2 + Cα-mCherry cells (τ = 2.4 ± 0.227 ns, n = 54 cells; P = 2.91×10^−5^) (Fig. 6E). Puncta fluorescence lifetime values in RIα^A99S^-GFP2 + Cα-mCherry cells were also similar to those recorded in puncta from cells only expressing WT RIα-GFP2 (τ = 2.52 ± 0.134 ns, n = 39 cells; ns, not significant vs RIα^A99S^-GFP2 + Cα-mCherry; P = 0.0066 vs. WT RIα-GFP2 + Cα-mCherry) (Fig. 6E). These live-cell observations were further corroborated by data showing that purified PKA-C (10% EGFP-tagged) localizes within liquid droplets when mixed with RIα and cAMP *in vitro* (Suppl. Fig 6F). Together, these data provide direct evidence for association between RIα and Cα within condensates and reveal that the pseudosubstrate IS plays a critical role in the formation of an active, non-canonical R:C complex.

Consistent with our previous study^21^, PKA-C not only localizes to RIα condensates but is also highly active within these phase-separated bodies, as detected by a green-red FRET-based PKA activity biosensor, FluoSTEP-AKAR^54^, in RIα-KO HEK293T cells co-expressing RIα tagged with the 11^th^ strand of sfGFP (RIα-FP11) (Fig. 6F, Suppl. Fig. 6G). We monitored the basal level of PKA activity within condensates by treating biosensor-expressing cells with the PKA inhibitor H89, followed by washout and application of Fsk/IBMX. PKA inhibition should promote dephosphorylation of the biosensor, leading to a decrease in the red-to-green (R/G) emission ratio, which is then normalized to the maximal R/G emission ratio produced by cAMP-induced re-phosphorylation of the biosensor by activated PKA. Compared with RIα-KO HEK293T cells re-expressing WT RIα-FP11, we observed significantly lower basal activity in cells expressing either RIα^A99S^-FP11 or WT RIα-FP11 plus the linker peptide (RIα(62-113)) that competes for PKA-C (Fig. 6F, Suppl. Fig. 6G), suggesting that PKA-C co-recruitment to RIα condensates via the non-canonical R:C complex is responsible for high basal PKA activity detected within RIα condensates.

PKA-C sequestration within RIα condensates may also impact global (e.g., cytosolic) PKA activity outside condensates. To determine whether cytosolic PKA activity is influenced by R:C interactions within RIα puncta, we used our cyan/yellow FRET-based PKA sensor AKAR4^55^ as a surrogate PKA substrate, which we expressed in RIα-KO HEK293T cells with or without mRuby2-tagged WT RIα or RIα^A99S^ co-expression. We monitored the level of basal PKA activity in the cytosol by examining the AKAR4 response in cells following treatment with H89 and Fsk/IBMX as above. Consistent with the enrichment of active PKA within condensates, we observed not only a 2.29-fold larger basal activity ((R_0_-R_min_)/ΔR_max_) but also a significantly higher starting Y/C ratio (R) in RIα-KO cells transfected with RIα^A99S^ ((R_0_-R_min_)/ΔR_max_ = 0.376 (0.300-0.453), n = 56 cells, P = 1.23×10^−5^ vs. RIα; R = 0.947 ± 0.128, n = 101 cells, P < 1×10^−15^ vs. WT) than in cells transfected with WT RIα ((R_0_-R_min_)/ΔR_max_ = 0.164 (0.109-0.189), n = 49 cells; R = 0.797 ± 0.095, n = 84 cells) (Fig. 6G, Suppl. Fig. 6H-J), reflecting elevated basal PKA activity. These results suggest that loss of IS-mediated PKA-C recruitment to RIα condensates increases basal PKA activity in the cytosol.

## DISCUSSION

Phase separation of the RIα PKA subunit to form cAMP-sequestering biomolecular condensates is emerging as a major driver of spatial compartmentation and signaling specificity in the cAMP/PKA pathway^21^. Here, we were able to identify key molecular determinants that control the ability of RIα to undergo LLPS. Two distinct interaction interfaces, the well-characterized D/D domain and the recently described N3A motif that abuts the CNB-A domain, were both required for RIα LLPS, as neither interface alone was sufficient to enable RIα condensate formation (Fig. 1 and 3). PKA-C release from the pseudosubstrate IS motif, which exposes the disordered linker joining the D/D and CNB-A domains, was similarly required for RIα LLPS. Mutations that promoted PKA-C release from the IS greatly enhanced condensate formation (Fig. 5). Conversely, incorporating CNB-domain mutations that locked PKA-C in place by breaking the cAMP-triggered allosteric switch, thereby preventing exposure of the disordered linker (Fig. 4), as well as disrupting the LLPS-promoting characteristics of the linker region by removing charged residues (Suppl Fig. 5E), abolished RIα LLPS. Our results complement the extensively documented roles of multivalency and intrinsic disorder in driving liquid-like condensate formation^18^. Notably, the positioning of the D/D and N3A sites on either side of the disordered linker is also reminiscent of the sticker and spacer model of condensate formation^47^. Whether these interaction sites can promote the requisite higher-order assemblies or if additional interactions are involved requires further investigation.

Both here (Fig. 6) and in our previous study^21^, we observed that fluorescently tagged PKA-C localizes within RIα phase-separated puncta in cells. The textbook view of PKA activation holds that the R:C complex fully dissociates upon cAMP binding^56–58^. Recent work seemed to overturn this model and argue that active RII:C holoenzymes remain intact during physiological cAMP elevations^14^. However, this mechanism was quickly refuted by a subsequent study^16^, leaving an unclear record of R:C complex dissociation. We found that PKA-C recruitment into RIα condensates is mediated by the IS motif, indicating a direct physical interaction, which is supported by the colocalization of purified Cα and RIα within RIα dropletes *in vitro* (Supplementary Fig. 6). Indeed, mutating the IS to promote PKA-C release, which strongly enhanced RIα LLPS (Fig. 5), largely eliminated the co-distribution of PKA-C into RIα condensates (Fig. 6). Similarly, expression of a peptide containing the IS motif successfully competed for PKA-C and prevented its recruitment to RIα condensates (Supplementary Fig. 6). Consistent with our previous data, PKA-C is catalytically active within RIα condensates (Fig. 6)^21^. Collectively, these results strongly suggest the existence of a bona fide non-canonical holoenzyme conformation in which cAMP-bound RIα and active PKA-C remain physically tethered but lack their mutual inhibitory behavior, allowing RIα to form condensates and PKA-C to phosphorylate substrates. Formation of this non-canonical complex may be facilitated by local RIα enrichment via LLPS, similar to a previous model of excess R subunit expression relative to PKA-C functioning to locally “trap” PKA-C through release and recapture^16^. This non-canonical R:C complex may therefore be limited to RI isoforms, considering that RII subunits have not been shown to phase separate and contain a true PKA substrate sequence within the IS, which we would expect to prevent PKA-C colocalization due to dissociation of the complex^53^. Indeed, such isoform-dependence was observed in early work examining the dynamics of dye-aggregation-induced fluorescent R:C puncta, potentially foreshadowing condensates^59^. RII-mediated PKA-C trapping may nevertheless come into play via local enrichment of R subunits by AKAP nanoclusters^60–62^. Interestingly, we found that AKAP docking to the D/D domain inhibited cytosolic RIα LLPS (Fig. 2), highlighting distinct but parallel mechanisms for spatially restraining PKA-C through molecular assembly. At the same time, RIα molecules recruited to membranes via AKAP binding may plausibly form membrane-proximal condensates or nanoclusters, as hinted by the uneven distribution of fluorescently tagged RIα at the plasma membrane (Figure 2C). Nevertheless, whereas AKAPs are known to direct PKA activity towards specific targets^29,59,63,64^, among other roles^9,10^, the function of non-canonical PKA-C recruitment to RIα condensates is unclear. One possible role may be in tuning cytosolic PKA activity, as suggested by our findings that disrupting the non-canonical R:C complex leads to increased basal phosphorylation by PKA (Fig. 6). Alternatively, RIα condensates may function as a mobile pool of PKA-C^65^ for nuclear and/or mitochondrial signaling. PKA activity may also play a direct role within RIα condensates. Futures studies will be aimed at exploring these possibilities.

In summary, the main advancements of this work include: 1) new molecular insights into the formation of RIα biomolecular condensates; 2) novel experimental evidence for the existence of a non-canonical PKA holoenzyme conformation within RIα condensates, which revises the classical model of PKA activation; and 3) demonstration of a novel role for RIα condensates in redistributing enzymatic activity to enable signaling specificity, showcasing the functional versatility of biomolecular condensates. Dysregulated cAMP/PKA signaling is linked to numerous diseases^52,63,66–74^, and we previously showed that loss of RIα LLPS can trigger tumorigenic phenotypes^21^. Loss of cAMP compartmentation results in aberrant cAMP elevations, which would likely impact multiple downstream cAMP effectors. However, our current findings reveal that RIα LLPS directly shapes PKA signaling, as exclusion of PKA-C from condensates leads to elevated cytosolic PKA activity. Pathological signaling may thus arise through loss of cAMP compartmentation, PKA recruitment, or a combination of both. Future studies will be aimed at dissecting the role of these processes in driving pathological signaling in diseases such as cancer and diabetes. Our results underscore the importance of LLPS as an organizer of the signaling architecture.

### Limitations of the study

This study provided clear evidence for the existence of a non-canonical PKA holoenzyme conformation in which cAMP-bound RIα and active PKA-C remain physically tethered but lack their mutual inhibitory activities. We showed that this non-canonical conformation is mediated by the disordered linker of RIα, and more specifically, the pseudosubstrate inhibitory sequence (IS) embedded within the linker. However, due to the limitations of the experimental approaches used here, we lack information on the structural differences between the canonical and non-canonical holoenzyme conformations, especially in the disordered linker region, and how these structural differences may lead to the striking differences in signaling activities discovered in this study. Structural approaches will be used in future studies to provide further insights at atomic resolution.

## ACKNOWLEDGEMENTS

The authors thank Dr. Friedrich Herberg (University of Kassel) for providing RIα-Rluc8 and GFP2-Cα constructs and assistance with BRET experiments. We are also grateful to Eric Griffis and Peng Guo from the UCSD Nikon Imaging Center for their training and assistance with the spinning-disk confocal microscope and the UCSD Neuroscience Microscopy Core for their training and assistance with the confocal microscope, which is supported by NIDNS P30NS047101. This work was supported by NIH R35 CA197622, NIH R01 DK073368 and a Fibrolamellar Cancer Foundation grant (to J.Z) and NIGMS GM102362 (to DW).

## AUTHOR CONTRIBUTIONS

J.C.H, S.S.T., and J.Z. conceived the project. J.C.H. and J.Z. designed experiments. J.C.H., E.H.P., Q.L., X.Z., D.R.Z, M.P., J.L.C.C., S.S., D.W., and S.M. performed experiments and analyzed the data. J.G.H.B., G.T., L.C., S.S. and H.N.L. provided critical reagents. S.S.T. and J.Z. supervised the project and coordinated experiments. J.C.H. generated final figures. J.C.H., S.M., and J.Z. wrote the manuscript.

## STAR ★ METHODS

### KEY RESOURCES TABLE

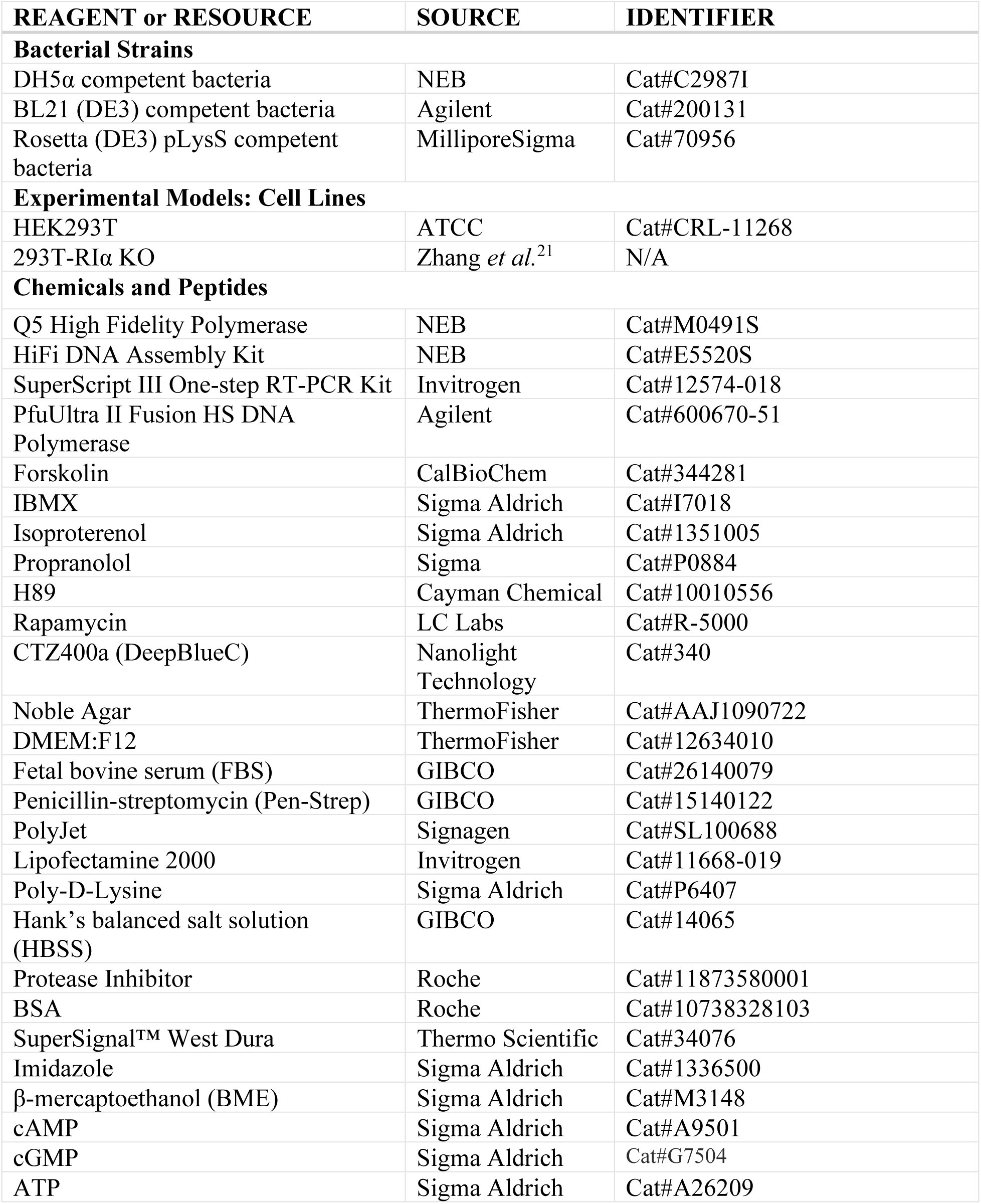
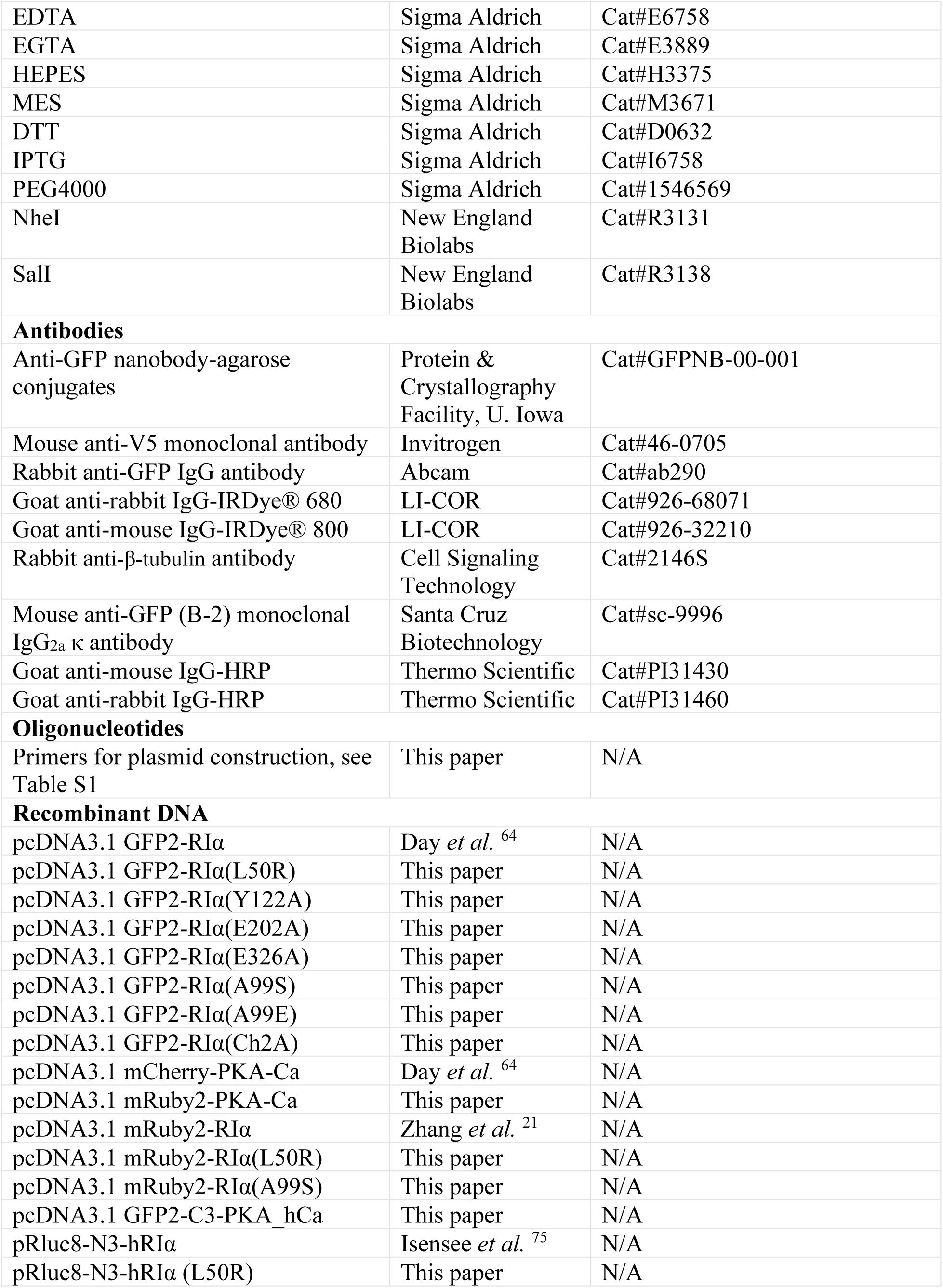
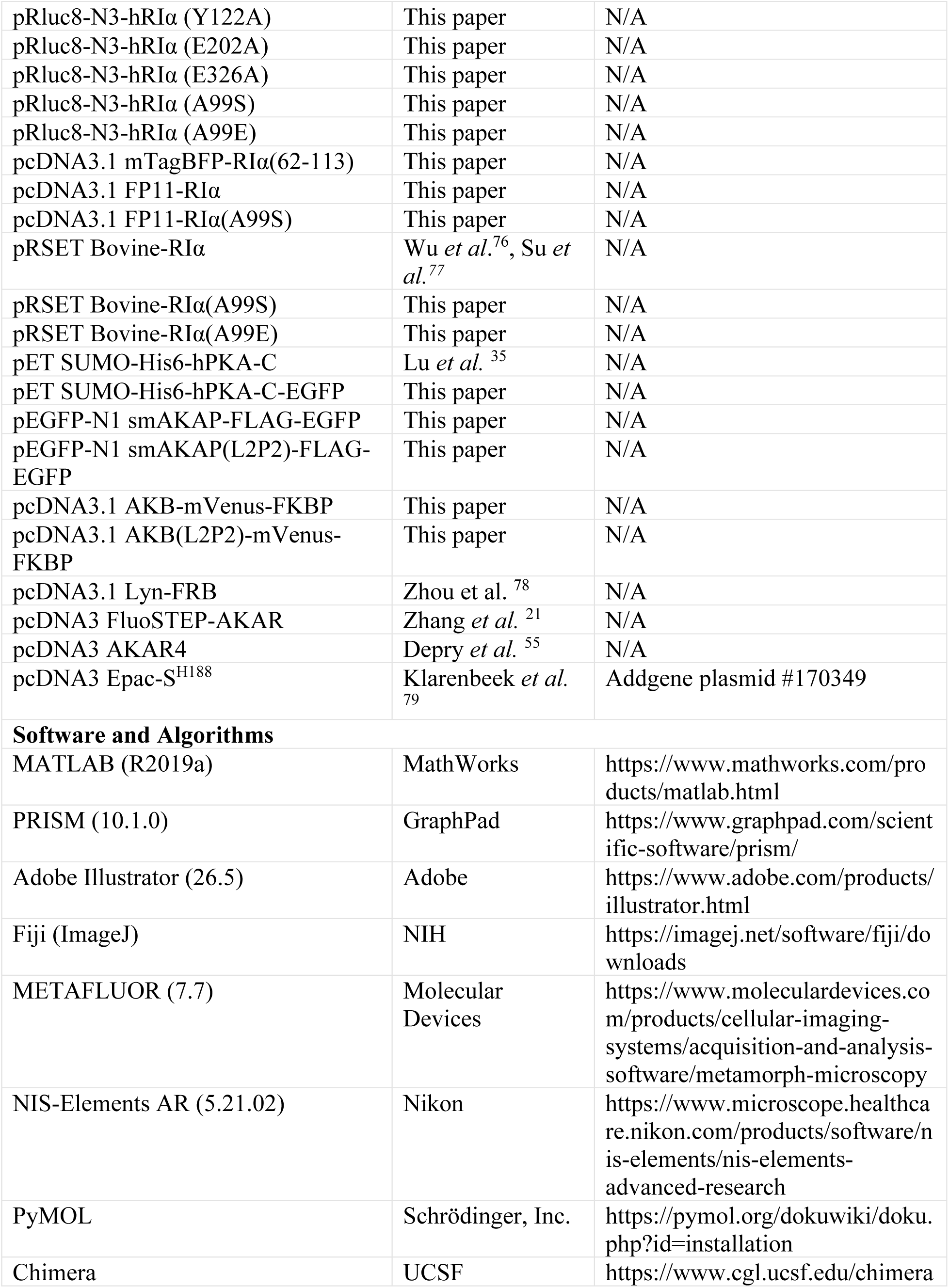

### RESOURCE AVAILABILITY

#### Lead contact

Further information and requests for resources and reagents should be directed to and will be fulfilled by the Lead Contact, Jin Zhang.

#### Materials availability

Plasmids generated in this study will be made available through Addgene and can be shared upon request.

### EXEPRIMENTAL MODEL AND SUBJECT DETAILS

#### Cell Culture and Transfection

HEK293T and RIα KO HEK293T cells were maintained in cell culture-treated plates with DMEM containing 1 g/L glucose, L-Glutamine, 110 mg/L Sodium Pyruvate with phenol red (GIBCO) and supplemented with 10% (v/v) fetal bovine serum (FBS, Sigma) and 1% (v/v) penicillin-streptomycin (Pen-Strep, Sigma-Aldrich). All cells were maintained in 5% CO_2_ humidified incubator at 37°C. Before transfection, cells were seeded on poly-D-lysine-coated 35-mm glass-bottom imaging dishes or 6-well plates and grown to 50-70% confluency. Cells were subsequently transfected using PolyJet (Signagen) and incubated an additional 16-24 hrs before imaging.

### METHOD DETAILS

#### Plasmid construction

All plasmids were generated in the pcDNA3.1 backbone (Invitrogen) unless specified otherwise. Primers used to generate plasmids are listed in Supplementary Table 2. All constructs were verified by Sanger sequencing (Genewiz, unless otherwise stated).

##### RIα mutant constructs

GFP2-RIα^64^, mRuby2-RIα^21^, pRLuc8-N3-RIα^75^ (gift from Dr. Friedrich Herberg, University of Kassel), and pRSET Bovine-RIα^76,77^ were described previously. All RIα point mutations were introduced via Gibson Assembly by cloning PCR products amplified from GFP2-RIα, mRuby2-RIα, RLuc8-N3-RIα, or Bovine-RIα using the NEBuilder Hi-Fi DNA Assembly Kit (New England Biolabs).

##### Fluorescent protein-tagged PKA-Cα constructs

mRuby2-PKA-Cα in pcDNA3.1 was generated via PCR amplification (Q5 High Fidelity Polymerase, New England Biolabs) of mRuby2 from mRuby2-RIα^21^ and PKA-Cα from mCherry-PKA-Cα, followed by Gibson assembly (NEBuilder Hi-Fi DNA Assembly Kit, New England Biolabs). pcDNA3.1 GFP2-C3-PKA_hC was generated via PCR amplification (Q5 High Fidelity Polymerase, New England Biolabs) of the pcDNA3.1 backbone from mCherry-PKA-Cα^64^ and GFP2-C3-PKA_hCa from the pGFP2-C3-PKA_hCa^80^, followed by Gibson Assembly (NEBuilder Hi-Fi DNA Assembly Kit, New England Biolabs). pET EGFP-SUMO-His6-hPKA-C was generated with PCR amplified pET SUMO-His6-hPKA-C backbone and GFP2 from RIα-GFP2. After these two fragments were assembled with Gibson Assembly (NEBuilder Hi-Fi DNA Assembly Kit, New England Biolabs), the S76T were introduced via Gibson Assembly by cloning PCR products amplified to make GFP2 into EGFP.

##### smAKAP-FLAG-EGFP and smAKAP(L2P2)-FLAG-EGFP

The cDNA of human smAKAP (c2orf88; NM_001042521.2) was cloned by RT-PCR from HEK293 cells using the SuperScript III one-step RT-PCR kit (Invitrogen). The forward primer included a NheI site, while reverse primers included a FLAG epitope tag as well as a SalI site (synthesized as two overlapping primers). After digestion, the cDNA was ligated into pEFGP-N1 (Clontech/Takara) digested with the same restriction sites to encode a smAKAP-FLAG-EGFP fusion protein. The PKA binding-deficient L65P, L70P (L2P2) mutant was generated according to the QuikChange protocol using PfuUltra II Fusion HS DNA polymerase and the supplied buffer (Agilent). All primers were purchased from IDT (Coralville, IA).

##### AKB-mVenus-FKBP and AKB(L2P2)-mVenus-FKBP

The A-kinase-binding (AKB) domains of WT (TVILEYAHRLSQDILCDALQQWAC) and L2P2-mutant (TVILEYAHR<u>P</u>SQDI<u>P</u>CDALQQWAC; L65P and L70P mutations underlined) smAKAP were PCR amplified from pEGFP-N1 smAKAP-FLAG-NES-EGFP and pEGFP-N1 smAKAP(L2P2)-FLAG-NES-EGFP, respectively. The resulting PCR fragment was inserted into a PCR-amplified pcDNA3.1 backbone containing mVenus and FKBP using Gibson assembly.

##### mTagBFP-RIα(62-113)

The RIα linker region (residues 62-113) and the pcDNA3.1 mTagBFP backbone were isolated from RIα(62-113)-GFP2 and RIα-mTagBFP^21^, respectively, using PCR amplification and re-assembled using Gibson assembly (NEBuilder Hi-Fi DNA Assembly Kit, New England Biolabs).

##### RIα-FP11

The 11^th^ strand of sfGFP (FP11; RDHMVLHEYVNAAGIT) was generated via PCR amplification of overlapping oligonucleotides. Additionally, the pcDNA3.1 RIα backbone was isolated and amplified from RIα-GFP2^21^ with PCR. The FP11 fragment was then inserted at the C-terminus of RIα via Gibson assembly (NEBuilder Hi-Fi DNA Assembly Kit, New England Biolabs).

#### Protein purification

##### pRSET Bovine-RIα

Bovine PKA RIα was purified as previously described^76,77^. WT and mutant bovine RIα constructs were transformed into *E. coli* BL21 (DE3) cells. All subsequent cultures were supplemented with 0.1mg/mL ampicillin. Starter cultures (3 mL) were grown from single colonies and used to inoculate large-scale (1 L) cultures, which were incubated over-night at 37 °C, 200 rpm, until reaching an OD600 of 0.6-1. Cultures were then induced with 0.5 mM IPTG. After ~16 h of expression at 16 °C, 150 rpm, cells were harvested and stored at −20 °C.

Cell pellets were resuspended in lysis buffer (20 mM MES, pH 6.5, 100 mM NaCl, 5 mM DTT, 2 mM EDTA, 2 mM EGTA, and an in-house protease inhibitor cocktail) and lysed with a microfluidizer. Total cell lysates were centrifuged for 1 h at 13,000 rpm, and the supernatant was ammonium precipitated. The resulting pellet was resuspended in lysis and applied to in-house-prepared cAMP resin. The resin mixture was incubated overnight at 4 °C. The resin was then washed using 6 column volumes of lysis buffer, followed by 6 column volumes of lysis buffer with 0.7 M NaCl and another 6 column volumes of lysis buffer, all at 4 °C. Bound protein was eluted using 3 × 10 mL of 40 mM cGMP elution buffer at room temperature. The elutions were combined, concentrated and loaded onto an S200 gel-filtration column equilibrated with 50 mM MES, pH 5.8, 200 mM NaCl, 5 mM DTT, 2 mM EDTA and 2 mM EGTA. Peak fractions were analyzed by SDS-PAGE, pooled and dialyzed into assay buffer.

##### pET SUMO-His6-hPKA-C

Human PKA Cα was purified as previously described^35^. Briefly, WT PKA Cα subunit with or without an EGFP tag in a pET-His6-SUMO vector was transformed into Rosetta (DE3) pLysS cells (MilliporeSigma). Transformed cells were then cultured, induced, and harvested as described above for bovine RIα.

Cell pellets were resuspended in buffer (20 mM Tris, pH 8, 300 mM NaCl, 5 mM β-mercaptoethanol (BME) and complete EDTA-free protease inhibitor cocktail (Sigma-Aldrich)) and lysed with a microfluidizer. Total cell lysate was centrifuged for 1 h at 13,000 rpm, and the supernatant applied to 5-20 mL of equilibrated Ni-NTA Agarose (Qiagen) resin. Following incubation at 4 °C overnight, 5-8 column volumes of 20 mM Tris pH 8, 300 mM NaCl and 2 mM BME were applied to wash the resin, followed by 20-50 mL of elution buffer (wash buffer plus 150 mM imidazole) to elute the bound C subunit. The eluate was then incubated for 1 h at room temperature with previously purified His-tagged Ulp1 (1:200 molar ratio) to cleave the His-Sumo tag and then dialyzed overnight in 4 L of wash buffer to remove imidazole. The next day, dialyzed protein solution was incubated with 5-20 mL of Ni Sepharose resin for 1 h at room temperature to bind the cleaved His-Sumo tag and any uncleaved protein. The flow-through was then collected and concentrated before loading onto an S200 gel-filtration column equilibrated with 20 mM MES, pH 6.5, 200 mM NaCl and 5mM DTT. Peak fractions were analyzed by SDS-PAGE, pooled and dialyzed into assay buffer.

#### *In Vitro* liquid droplet assay

All liquid droplet formation assays were performed in 150 mM KCl, 5 mM MgCl^2^, 20 mM HEPES, pH 7.0, 1 mM EGTA, 1 mM DTT, 0.5 mM ATP, cAMP (as indicated), cAMP (as indicated), and Polyethylene Glycol 4000 as specified. After the protein concentration was determined using Thermo Scientific Pierce BCA Protein Assay kit (Thermo Fisher Scientific), purified proteins were incubated at different stoichiometries and at various concentrations at room temperature for 1 hr in glass-bottom, black-walled 96-well plates and imaged under DIC and/or fluorescence microscopy.

#### Immunoblotting and co-immunoprecipitation

##### RIα^L50R^-GFP2 expression verification

HEK293T RIα KO cells were plated in a 6-well plate and transfected with 1 µg RIα-GFP2 and RIα^L50R^-GFP2 using PolyJet (Signagen) according to manufacturer’s protocol 24 hrs later. The next day, cells were rinsed twice with phosphate-buffered saline (PBS) and harvested. The harvested cells were lysed in radioimmunoprecipitation assay RIPA buffer (10 mM Tris-Cl, pH 8.0, 0.1% sodium deoxycholate, 0.5 mM EGTA, 1 mM EDTA, 0.1% SDS, 1% Triton X-100, 140 mM NaCl) containing protease inhibitor (Roche Applied Science) for 15 min on ice. After centrifugation (12000RPM, 4°C for 15min), the supernatant was collected, and protein quantification was performed using a Thermo Scientific Pierce BCA Protein Assay kit (Thermo Fisher Scientific). 20 µg of protein sample per well was loaded on 4–20% Mini-PROTEAN® TGX™ Precast Protein Gels SDS–PAGE gels (BioRad) and electrophoresed for 1 h and 25 min. The separated proteins on the gel were transferred to 0.45 µm PVDF membrane (Millipore). After blocking (5% BSA in 0.1% PBS-Tween (PBST) for 30 min), the membranes were incubated in the diluted primary antibodies (beta-tubulin: 1:5000, GFP: 1:200) at 4°C for overnight. Membranes were washed 4 times for 5 minutes each with 0.1% TBS-Tween (with TBST), then incubated with a diluted secondary antibody (1:5000) for 1 hour at room temperature. The chemiluminescent HRP substrate (SuperSignal™ West Dura, Thermo Scientific) was added to the membrane and then imaged using ChemiDoc™ Gel Imaging System (BioRad). Immunoblot data were quantified using ImageJ.

##### smAKAP^L2P2^ mutation verification

HEK293 RIα KO cells were plated at 600,000 cells/well in 6-well plates and transfected the next day with 1 μg each of RIα-V5-LgBiT and smAKAP-GFP (WT or L2P2 mutant) using Lipofectamine 2000 according to the manufacturer’s protocol. The following day, cells were lysed and subjected to co-immunoprecipitation as previously described^81^ using anti-GFP nanobodies conjugated to agarose beads (Protein & Crystallography Facility, U. Iowa). Protein complexes were separated by SDS-PAGE, transferred to nitrocellulose^81^, and probed sequentially with primary antibodies directed against GFP (Abcam) and the V5 epitope (Invitrogen) for 1 h each. Following 1 h incubation with appropriate IRDye-680- and IRDye-800-coupled secondary antibodies and additional washes, blots were imaged on a LI-COR Odyssey infrared scanner. GFP and V5 blots of the same IP lanes are shown.

#### Bioluminescence resonance energy transfer (BRET) assays

White-walled, clear-bottom 96-well assay plates (Costar) were coated with 0.1 mg ml^−1^ of poly-d-lysine (PDL) in Dulbecco’s phosphate-buffered saline (pH 7-7.3) for 24 h. Cells were transfected with PolyJet in 6-well plates (Costar) for 24 h and then seeded at 5 × 10^4^ HEK293T cells per well in the PDL-coated 96-well assay plates.

Luminescence intensities were recorded on a Spark 20M fluorescence microplate reader using SparkControl Magellan 1.2 software (TECAN). Cells were washed twice with HBSS and then placed in 100 µL of HBSS containing 5 µM CTZ400a (DeepBlueC, Nanolight Technology) immediately prior to recording for 8 cycles to obtain a baseline reading. To monitor R:C complex dissociation, wells received an additional 100 µL of CTZ400a-HBSS solution without (control) or with 100 µM Fsk (Calbiochem; final concentration 50 µM) and were read for an additional 15 cycles. For R subunit dimerization assays, wells were read for 10 cycles after receiving only 100 µL of HBSS containing 5 µM CTZ400a. Intensities were recorded from each well using the monochromator (400 ± 40 nm and 540 ± 35 nm) with a gain of 25 and a 1-sec integration time. Entire plate was read one wavelength at a time, resulting in a time interval of ~ 2 min per cycle.

#### Time-lapse fluorescence imaging

Cells were washed twice with HBSS and then imaged in HBSS in the dark at room temperature. Forskolin (Fsk, Calbiochem), 3-isobutyl-1-methylxanthine (IBMX, Sigma), Isoproterenol (Iso, Sigma Aldrich), Propranolol (Prop, Sigma), Rapamycin (Rapa, LC Labs), and H89 (Cayman Chemical) were added as indicated.

##### Widefield epifluorescence imaging

Puncta formation assays were imaged on a Zeiss Axiovert 200M microscope (Carl Zeiss) equipped with a 40×/1.3 NA oil-immersion objective, an ORCA-Flash4.0LT digital CMOS camera (Hamamatsu) and controlled by METAFLUOR 7.7 software (Molecular Devices). GFP intensity was imaged using a 480DF30 excitation filter, a 505DRLP dichroic mirror, and a 535DF45 emission filter. All filter sets were alternated using a Lambda 10-2 filter changer (Sutter Instruments). Exposure times for each channel were 500 ms, and images were acquired every 30 s.

AKAP colocalization, AKB translocation, puncta reversibility experiments, additional puncta formation assays, and Epac-S^H188^, FluoSTEP-AKAR and AKAR4 biosensor imaging experiments were performed on a Ziess AxioObserver Z1 microscope (Carl Zeiss) with equipped with Definite Focus system (Carl Zeiss), a 40×/1.4 NA oil-immersion objective, and a Photometrics Evolve EMCCD (Photometrics) and controlled by METAFLUOR 7.7 software (Molecular Devices). GFP intensities were imaged using a 480DF30 excitation filter, a 505DCXR dichroic mirror, and a 535DF45 emission filter. RFP intensities were imaged using a 555DF25 excitation filter, a ZT568RDC dichroic mirror, and a 650DF100 emission filter. BFP intensities were imaged using 405DF40 excitation filter, a 450DCXRU dichroic mirror, and 475DF40 emission filter. Dual red/green emission ratio imaging was performed using a 555DF25 excitation filter, a ZT568RDC, and two emission filters (650DF100 for RFP and 535DF45 for GFP). Dual cyan/yellow emission ratio imaging was performed using a 420DF20 excitation filter, a 450DCXRU dichroic mirror and two emission filters (475DF40 for CFP and 535DF25 for YFP). All filter sets were alternated using a Lambda 10-3 filter changer (Sutter Instruments). Exposure times for each channel were 500 ms, except 50 ms for YFP direct, with electron multiplying gain set to 50, and images were acquired every 30 s.

*In vitro* liquid droplet assay experiments were imaged on a Zeiss AxioObserver Z7 microscope (Carl Zeiss) equipped with a ×40/1.3 NA objective and a Photometrics Prime 95B sCMOS camera (Photometrics), controlled by the METAFLUOR v.7.7 software (Molecular Devices). DIC images were acquired with a HXP 120V lamp. EGFP intensities were imaged using a480DF20 excitation filter, T505LPXR dichroic mirror and 535DF50 emission filter. Filter sets were alternated using an LEP MAC6000 control module (Ludl Electronic Products). Exposure times were 100 ms for DIC images and 50 ms for EGFP images.

##### Spinning-disk confocal imaging

Additional puncta formation assays and colocalization assays were imaged on a Nikon Ti2 microscope (Nikon) equipped with a Yokogawa W1 confocal scanhead (Yokogaw), an Opti Microscan FRAP unit (Bruker Nano Inc, integrated by Nikon), a six-line (405, 445, 488, 515, 561, and 640 nm) LUN-F-XL laser engine with bandpass and long-pass filters (450/50 and 525/50) (Chroma), an Apo TIRF 100×/1.49 NA objective, and a Prime95B sCMOS camera (Photometrics) and operated with NIS Elements software (Nikon). Z-stacks with 0.2 µm step were obtained sequentially in each channel with 200 ms excitation at 20% laser power (405, 488, and 561 nm) every 1-2 min as indicated.

##### Fluorescence recovery after photobleaching (FRAP)

HEK293T cells were imaged on the spinning-disk confocal microscope described above. Circular regions of interest were drawn over whole puncta (approximately 0.30 μm-1.00 μm radius) or similarly sized regions of bulk cytosol for photobleaching. Each experiment contained a minimum of 10 images at 0.25-0.5-sec intervals with 200 ms excitation using the 488 nm laser line at 5-20% power, then puncta or bulk cytosol regions of interest were bleached once using the 405 nm laser line for 500 ms at 75-100% power and recovery monitored every 0.25-0.5 sec for a total of 1-2 min.

##### Fluorescence lifetime imaging microscopy (FLIM)

HEK293T cells were imaged using a Leica SP8 FALCON confocal microscope equipped with a 63×/X NA oil-immersion objective, a white-light laser, and a Lecia HyD SMD detector. The laser was tuned to 502-532 nm for excitation of RIα-EGFP and tuned to 582-611 nm for excitation of Cα-mCherry variants and pulsed at a frequency of 80 MHz. Images were acquired before (Time 0) and 15 min after (Time 15) Fsk/IBMX addition, collecting 1,000 photons per pixel.

### QUANTIFICATION AND STATISTICAL ANALYSIS

#### BRET biosensor analysis

Emission ratios were calculated as 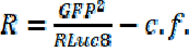 or each condition with the control factor 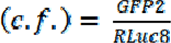 when only RLuc8 is present. Fsk-induced ratio change was calculated as 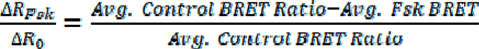. Graphs were plotted using Prism 10 (GraphPad).

#### FRET biosensor analysis

Raw fluorescence images were corrected by subtracting the background intensity from an ROI without cells. The background-subtracted emission ratios of biosensor-expressing cells were used to calculate the emission ratios (C/Y, Y/C, or R/G) at each time point (R). For some time-courses, the emission ratios (R) were normalized to the emission ratio at time t = 0 (R/R_0_), which is defined as the time-point immediately before drug addition. The ratio change after drug stimulation (ΔR/R_0_) was calculated as (*R_drug_* − *R*_0_)/*R*_0_ where R_drug_ is defined as the mean R value across all time points after the response has reached a plateau following stimulation. The normalized basal ratio (t = 0) is calculated as *R*_0_ − *R_min_*/Δ*R_max_* where R_min_ is defined as the minimum value after inhibitor addition and ΔR_max_ is defined as the maximum value of *R*_0_ − *R_min_* after stimulation. Graphs were plotted using Prism 10 (GraphPad).

#### Cellular puncta quantification

Raw fluorescence images from each channel were compared to ensure expression of each construct in cells selected for analysis. Brightness and contrast were adjusted for each cell using Fiji (ImageJ) to ensure optimal puncta counting.

#### FRAP analysis

Intensity counts for puncta and unbleached reference puncta (one punctum and one reference punctum in the same cell) were background subtracted using a nearby region without cells and normalized to the pre-bleach average intensity. Bleached puncta were normalized to the unbleached reference puncta and then to the bleached punctum’s minimum and maximum intensities to determine the percent recovery. The percent recovery curves were fitted to an exponential recovery curve using the ImageJ. Samples with bleached or reference puncta that moved out of the z-plane during experiments were truncated (beyond 20-30 sec) or excluded from analysis. Following curve fitting, samples with curve y-intercepts ≥ 0.15 were excluded from further analysis since intensity values were normalized to 0 at t = 0, the time of photobleaching. Half-times for each sample were derived from the ImageJ curve fitting tool using 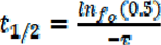 where τ = b in the exponential recovery equation *y* = *a*(1 − *e*)*^bx^* + *c*. Graphs were plotted using Prism 10 (GraphPad).

#### FLIM analysis

One region of interest (ROI) corresponding to an RIα condensate and one ROI corresponding to the cytosol were selected per cell. Amplitude-weighted mean fluorescence lifetimes were calculated in LAS X (Lecia) software by fitting a bi-exponential decay model to the decay of each ROI. Graphs were plotted using Prism 10 (GraphPad).

#### Partition coefficient

Integrated intensities of whole-cell and puncta ROIs were measured from 3 confocal z-slices (2-µm apart) using Fiji (ImageJ) to calculate the partition coefficient of tagged protein into puncta vs the cytosol (Σ*I_puncta_*/Σ*I_cell_*). These partition coefficients were plotted with respect to the ratio of the integrated whole-cell intensities (Σ*I_cell_*) of the RIα and Cα channels after Fsk/IBMX stimulation (+) to determine the relative expression dependencies of the partition coefficients. Graphs were plotted using Prism 10 (GraphPad).

#### Colocalization analysis

Line-scans were performed in Fiji (ImageJ) to determine colocalization. Emission intensities (I) were normalized by dividing the intensity at each position by the maximum intensity (I_max_) for reported line-scans (I/I_max_). Pearson’s Coefficient per RIα Puncta was calculated using NIS-Elements software. Percent colocalization was defined as the number of Cα-containing (i.e., RFP-positive) RIα puncta divided by the total number of RIα puncta per cell. Graphs were plotted using Prism 10 (GraphPad).

#### Statistics and reproducibility

Statistical analyses were performed in GraphPad Prism 9 (GraphPad). For normal data sets, pairwise comparisons were performed using unpaired Student’s t-tests, with Welch’s correction equal variances applied where indicated. Comparisons between more than two groups were performed using ordinary one-way analysis of variance (ANOVA) or Welch’s ANOVA, followed by the indicated multiple comparison test. For non-Gaussian (i.e., puncta quantification) data, pairwise comparisons before and after stimulation comparisons were performed using paired Wilcoxon signed-rank tests and comparisons to WT were performed using unpaired Kolmogorov-Smirnov tests. If n > 500, we used Cohen’s d (https://www.socscistatistics.com/effectsize/default3.aspx) to calculate the effect size. Statistical significance was set at P < 0.05. Throughout the paper, data sets combined at least three independent experiments, except puncta per cell data, which combined at least two independent experiments. Line-scans in Figures 2A, C-D, and 5H and Supplementary Figure 6C represented line-scans from at least ten cells.

