## Supplemental Figures and Tables for "Molecular Determinants and Signaling Effects of PKA RIα Phase Separation"

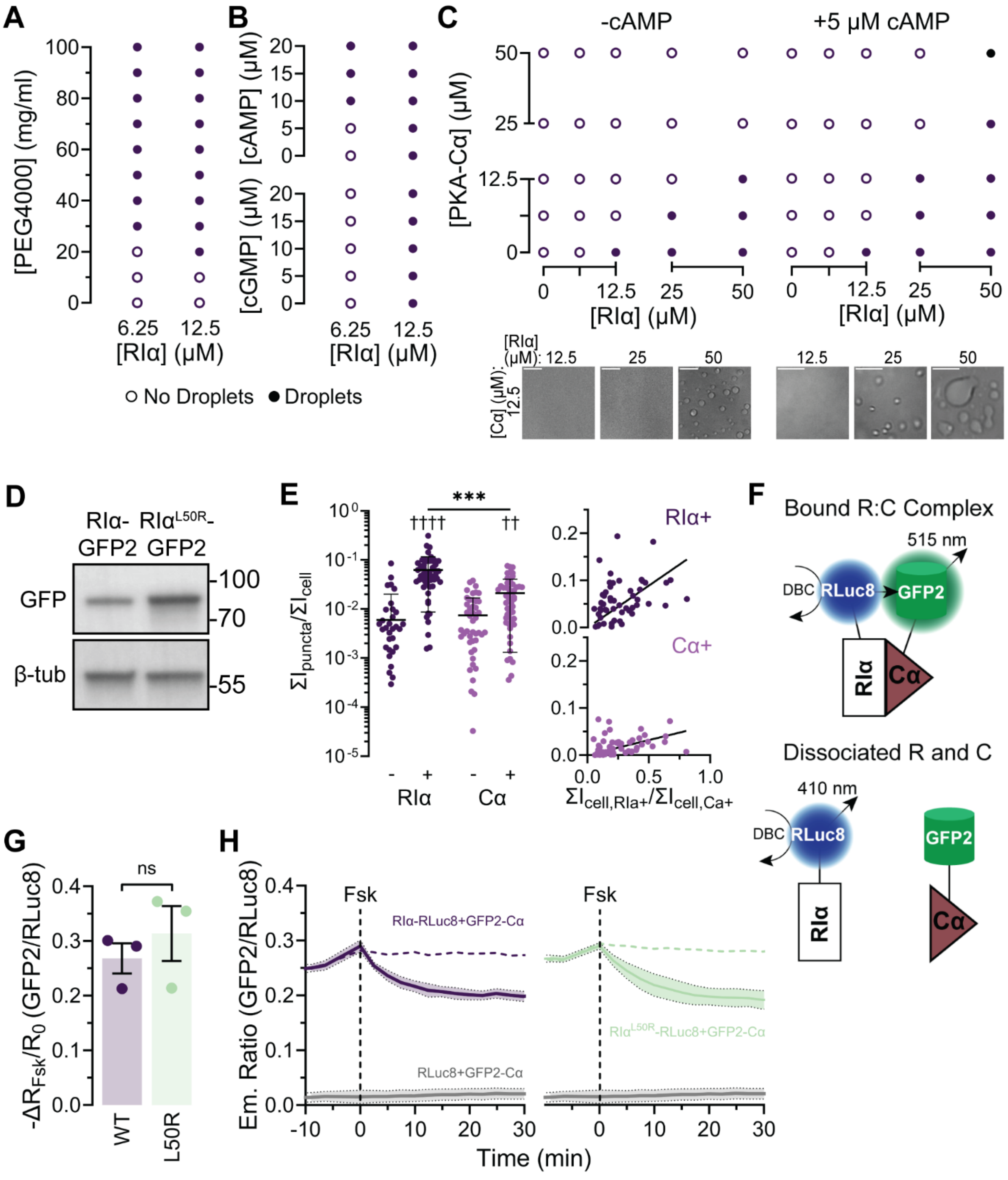


**Supplementary Figure 1. RIα LLPS and R:C dynamics of L50R mutant RIα. (A-C)** *In vitro* phase diagram of RIα vs PEG 4000 **(A)**, cAMP or cGMP **(B)** or Cα **(C)** with (right) or without (left) 5 μM cAMP. Representative images showing phase-separated liquid droplets with or without cAMP (bottom). Scale bars: 10 μm. **(E)** Western Blot of RIα-GFP2 and RIαL50R-GFP2 expressed in HEK293T RIα-KO cells using anti-GFP and anti-β-tubulin antibodies. Representative of 2 independent experiments. **(F)** Partition coefficient of puncta in HEK293T cells expressing RIα-GFP2 (RIα) and Cα-mRuby2 (Cα) before (-) and after (+) Fsk/IBMX stimulation (left) and partition coefficient vs. ratio of cellular of RIα expression relative to Cα after Fsk/IBMX stimulation (right). Error bars indicate mean ± SD. ***P = 0.000195 (+/RIα vs +/Cα), ††††P $< 1.0 \times 10^{-15}$ (-/RIα vs +/RIα) and ††P = 0.00125 (-/Cα vs +/Cα); Kruskal-Wallis test followed by Dunn's multiple comparisons test. **(G)** Schematic of BRET2 assay for monitoring R:C complex dissociation. **(H)** Summary of the Fsk-simulated change in the GFP2/RLuc8 emission ratios. n = 3 independent experiments. ns, not significant; unpaired, two-tailed Student's t-test. Bar graphs show mean ± SD. **(I)** Average time-course GFP2/RLuc8 emission ratios in HEK293T cells co-expressing GFP2-Cα plus RLuc8-fused to either WT RIα (left) or $RI\alpha^{L50R}$ (right). Dotted lines indicate data from vehicle-control wells. GFP2-Cα was co-overexpressed with RLuc8 alone as a negative control (solid gray curves). Curves are plotted as means (solid lines) ± SD (shading).

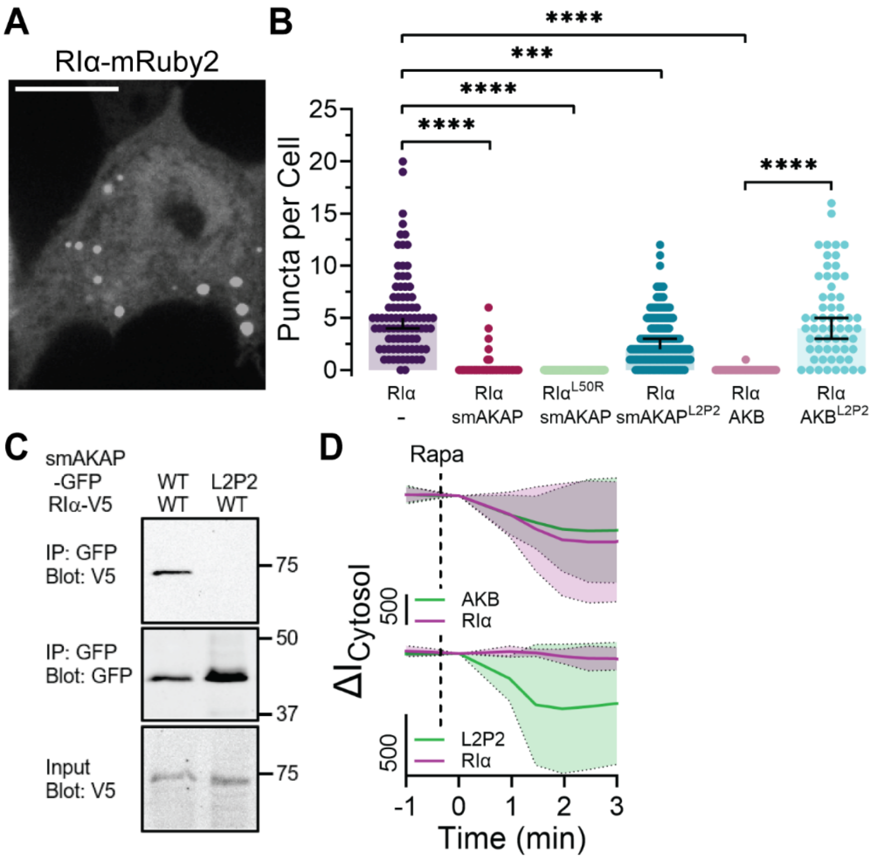


**Supplementary Figure 2. smAKAP binding inhibits RIα puncta formation. (A)** Representative maximum intensity projection from a confocal z-stack of a HEK293T cell overexpressing RIα-mRuby2. Scale bar, 10 µm. **(B)** Summary quantification of puncta number per cell in HEK293T cells expressing RIα-mRuby2 or $RIα^{L50R}$-mRuby2 alone or with either GFP-tagged WT or L2P2-mutated smAKAP or AKB. n = 87 (RIα), 312 (RIα + smAKAP), 403 ($RIα^{L50R}$ + smAKAP), 186 (RIα + $smAKAP^{L2P2}$), 59 (RIα + AKB), and 60 (RIα + $AKB^{L2P2}$) cells from 3 experiments each. Error bars indicate median ± 95% CI. ****$P < 1 \times 10^{-15}$ (RIα + smAKAP vs RIα-), ****$P < 1 \times 10^{-15}$ ($RIα^{L50R}$ + smAKAP vs RIα-), ***P = 0.000919 (RIα + $smAKAP^{L2P2}$ vs RIα-), ****$P < 1 \times 10^{-15}$ (RIα + AKB vs RIα-), and ****$P < 1 \times 10^{-15}$ (RIα + $AKB^{L2P2}$ vs RIα + AKB); Kruskal-Wallis test followed by Dunn's multiple comparisons test. **(C)** Representative western blots of co-immunoprecipitation from lysates of HEK293T cells co-expressing RIα-V5 plus either smAKAP-GFP or $smAKAP^{L2P2}$-GFP. **(D)** Average time-course of the rapamycin (Rapa, 1 µM)-included change in the cytosolic fluorescence intensity ($ΔI_{Cytosol}$) of

HEK293T cells co-expressing Lyn-FRB plus RIα-mRuby2 (purple curves) and either WT (upper) or L2P2-mutant (lower) AKB-mVenus-FKBP (green curves).

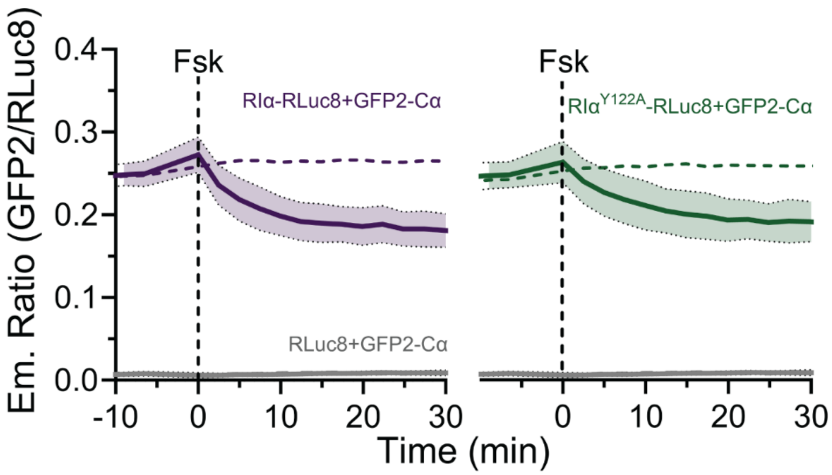


**Supplementary Figure 3. R:C dynamics with Y122A mutant RIα.** Average time-courses of the GFP2/RLuc8 emission ratio in HEK293T cells co-expressing GFP2-Cα plus RLuc8 fused to either RIα or RIα$^{Y122A}$ and stimulated with Fsk. Dotted lines represent data from vehicle-control wells. GFP2-Cα was co-overexpressed with RLuc8 alone as a negative control (solid gray curves). Data are from 3 independent experiments. Curves are plotted as means (solid lines) ± SD (shading).

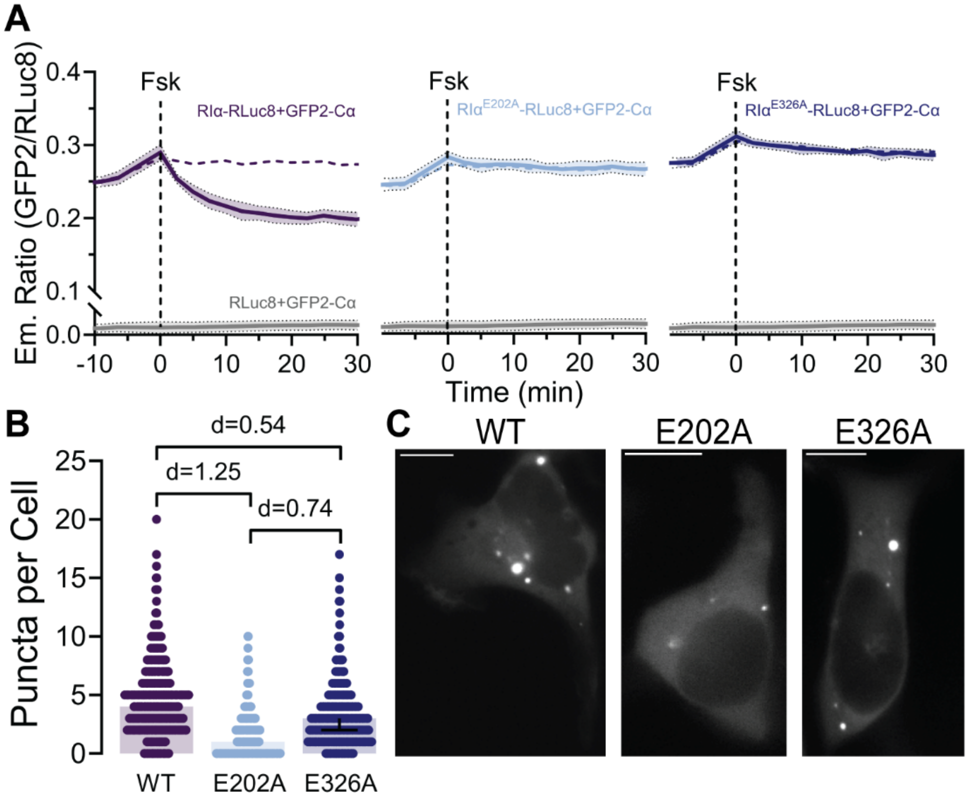


**Supplementary Figure 4. R:C dynamics with RIα CNB mutations. (A)** Average time-courses of the GFP2/RLuc8 emission ratio in HEK293T cells co-expressing GFP2-Cα plus RLuc8 fused to RIα (left), RIα$^{E202A}$ (middle), or RIα$^{E326A}$ (right) and stimulated with Fsk. Dotted lines indicate data from vehicle-control wells. GFP2-Cα was co-overexpressed with RLuc8 alone as a negative control (solid gray curves). Data are from 3 independent experiments. Curves are plotted as means (solid lines) ± SD (shading). **(B)** Summary of puncta number per cell in HEK293T cells expressing RIα-GFP2 (WT; n = 686 cells), RIα$^{E202A}$-GFP2 (E202A; n = 730 cells), and RIα$^{E326A}$-GFP2 (E326A; n = 708 cells). Effect size was calculated using Cohen's d. Error bars indicate median ± 95% CI. Data are from 2 experiments. **(C)** Representative images RIα-GFP2 (left), RIα$^{E202A}$-GFP2 (middle), and RIα$^{E326A}$-GFP2 (right) overexpressed in HEK293T cells. Scale bars: 10 µm.

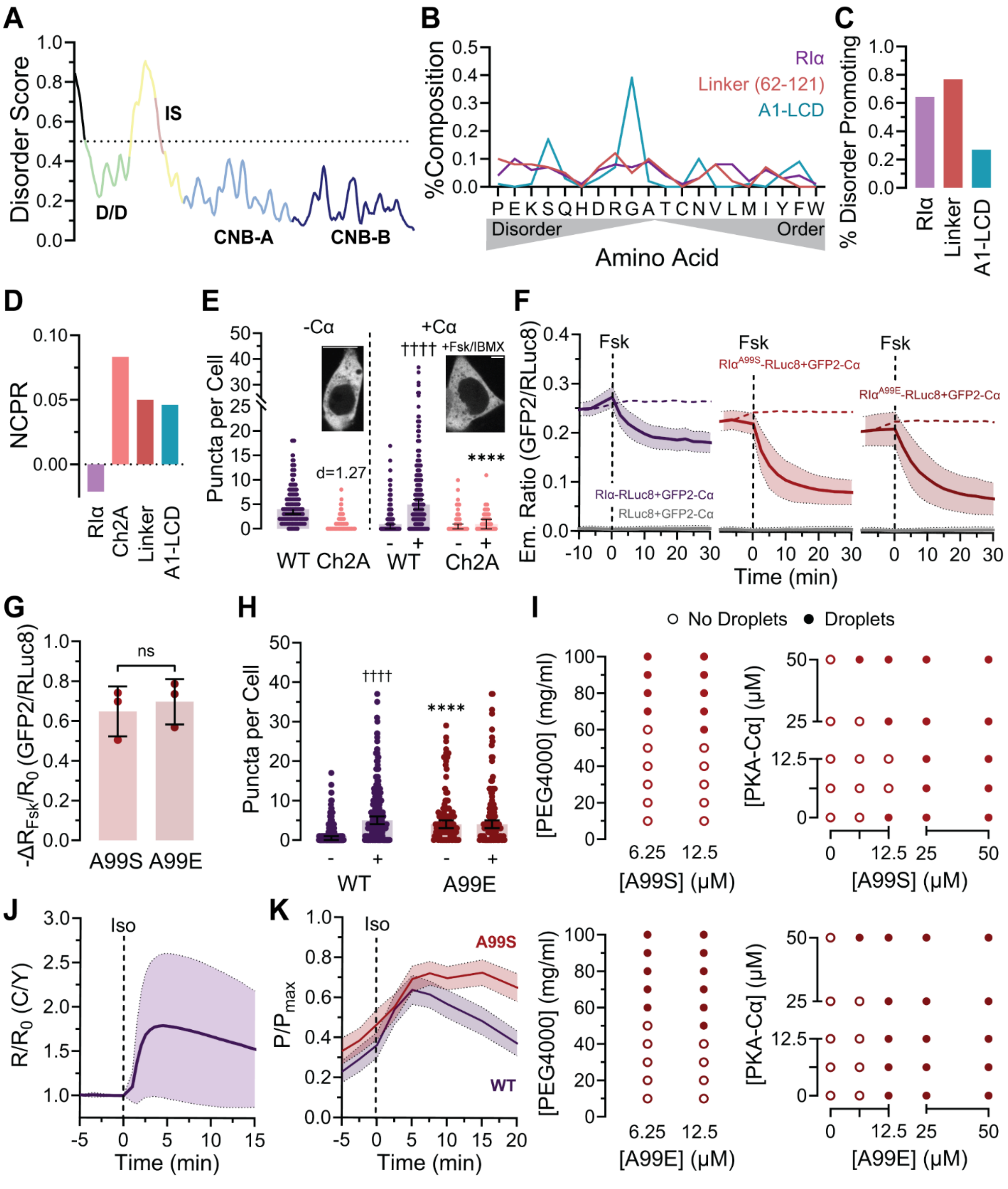


**Supplementary Figure 5. Further characterization of RIα IS mutations. (A)** IUPred3 long disorder analysis score of RIα amino acid sequence. **(B)** Percent composition of each amino acid in full-length RIα (RIα), residues 62-121 of RIα (Linker), and hnRNPA1 low-complexity domain

(A1-LCD) as a reference, organized from disorder to order promoting residues. Calculated from amino acid sequences using CIDER. **(C)** Percent of disorder promoting residues (P, E, K, S and Q) in RIα, Linker, and A1-LCD. Calculated from amino acid sequences using CIDER. **(D)** Net charge per residue (NCPR) of RIα, linker with charged residues (K, R, D, and E) mutated to Ala (Ch2A), Linker, and A1-LCD. Calculated from amino acid sequences using CIDER. **(E)** Summary of puncta number in HEK293T cells expressing RIα-GFP2 or RIα$^{Ch2A}$-GFP2, with charged residues (K, R, D, and E) in the linker region mutated to Ala, without (left; WT: n = 319 cells; Ch2A: n = 512 cells; 3 experiments) or with Cα-mCherry (right; WT: n = 255 cells from 6 experiments; Ch2A: n = 60 cells from 3 experiments). Inset shows representative confocal fluorescence images of HEK293T cells expressing RIα$^{Ch2A}$-GFP2 with or without Cα. Scale bars: 10 µm. Error bars indicate median ± 95% CI. ††††$P < 1 \times 10^{-15}$ (WT+ vs WT-); paired Wilcoxon rank sum test and ****$P = 9.08 \times 10^{-10}$ (Ch2A+ vs WT+); unpaired Komogorov-Smirnov test. **(F)** Average time-courses of the GFP2/RLuc8 emission ratio in HEK293T cells co-expressing GFP2-Cα plus RLuc8 fused to RIα (left), RIα$^{A99S}$ (middle), or RIα$^{A99E}$ (right) and stimulated with Fsk. Dotted lines represent data from vehicle-control wells. GFP2-Cα was co-overexpressed with RLuc8 alone as a negative control (solid gray curves). Curves are plotted as means (solid lines) ± SD (shading). Data are from 3 independent experiments. **(G)** Fsk-stimulated change in the GFP2/RLuc8 emission ratio in HEK293T cells co-expressing GFP2-Cα plus RIα$^{A99S}$-RLuc8 (A99S; n = 3 experiments) or RIα$^{A99E}$-RLuc8 (A99E; n = 3 experiments). ns, not significant; unpaired, two-tailed Student's t-test. Error bars indicate mean ± SD. **(H)** Summary of puncta number per cell before (-) and after (+) Fsk/IBMX stimulation in HEK293T cells co-expressing Cα-mCherry plus either RIα-GFP2 (reproduced from E) or RIα$^{A99E}$-GFP2 (A99E; n = 107 cells). ****$P = 4.37 \times 10^{-8}$ (A99E- vs WT-), unpaired Komogorov-Smirnov test; ††††$P < 1 \times 10^{-15}$

(WT+ vs WT-); paired Wilcoxon rank sum test. Error bars indicate median ± 95% CI. Data are from 3 experiments. **(I)** *In vitro* phase diagram of purified RIα$^{A99S}$ (top) and RIα$^{A99E}$ (bottom) vs. PEG 4000 (left) and Cα (right). **(J)** Average C/Y emission ratio time-course in HEK293T cells expressing the cAMP biosensor Epac-S$^{H188}$ upon treatment with isoproterenol (Iso; 10 μM). **(K)** Average time-course showing the max-normalize number of RIα puncta ($P/P_{max}$) in HEK293T cells co-expressing Cα-mCherry plus either RIα-GFP2 (purple curve) or RIα$^{A99S}$-GFP2 (red curve) with Iso treatment. Curves in **J** and **K** are plotted as means (solid lines) ± SD (shading).

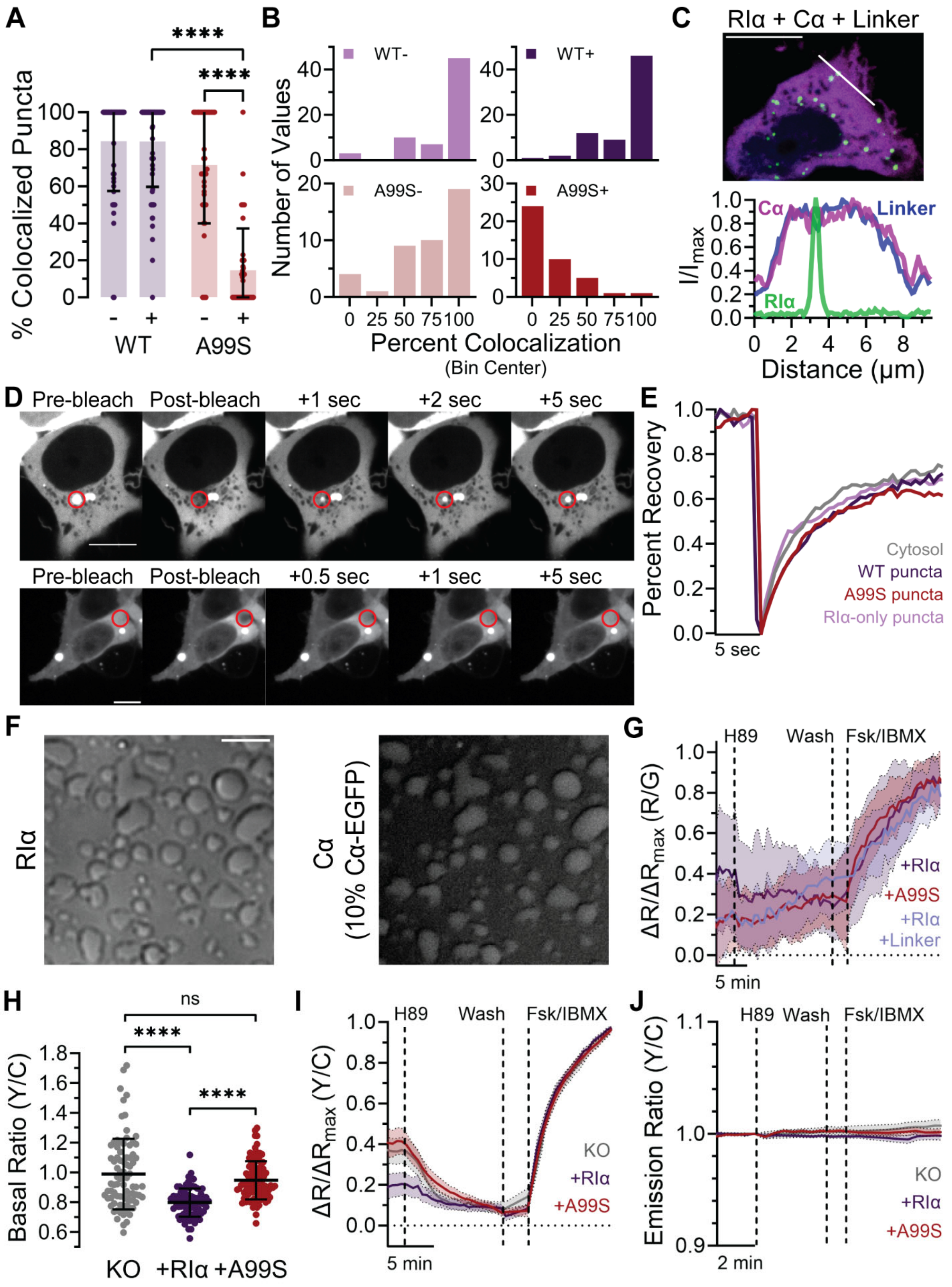
A
% Colocalized Puncta
****
****
WT
A99S
B
Number of Values
WT-
WT+
A99S-
A99S+
Percent Colocalization
(Bin Center)
C
RIα + Cα + Linker
Cα
Linker
RIα
I/I max
Distance (μm)
D
Pre-bleach
Post-bleach
+1 sec
+2 sec
+5 sec
+0.5 sec
E
Percent Recovery
Cytosol
WT puncta
A99S puncta
RIα-only puncta
5 sec
F
RIα
Cα
(10% Cα-EGFP)
G
H89
Wash
Fsk/IBMX
ΔR/ΔRmax (R/G)
+RIα
+A99S
+RIα
+Linker
5 min
H
ns
****
****
Basal Ratio (Y/C)
KO
+RIα
+A99S
I
H89
Wash
Fsk/IBMX
ΔR/ΔRmax (Y/C)
KO
+RIα
+A99S
5 min
J
H89
Wash
Fsk/IBMX
Emission Ratio (Y/C)
KO
+RIα
+A99S
2 min

**Supplementary Figure 6. Additional analysis of the non-canonical R:C complex. (A)** Summary of the percent colocalization of Cα-mRuby2 puncta with RIα-GFP2 (WT; n = 65 before and n = 70 after stimulation) or RIα$^{A99S}$-GFP2 (A99S; n = 43 before and n = 41 after stimulation) puncta before (-) and after (+) Fsk stimulation. ****P = $8 \times 10^{-15}$ (A99S+ vs WT+) and ****P = $4.6 \times 10^{-14}$ (A99S+ vs A99S-); two-way ANOVA followed by Tukey's multiple-comparisons test. Error bars indicate mean ± S.D. **(B)** Probability distributions showing the percent colocalization between Cα-mRuby2 puncta and RIα-GFP2 or RIα$^{A99S}$-GFP2 puncta before (-) and after (+) Fsk stimulation (bin width: 25). **(C)** Representative confocal images **(top)** and line-intensity profile of the indicated region in HEK293T cells **(bottom)** when RIα(62-113)-mTagBFP (Linker) is expressed with RIα-GFP2 and Cα-mRuby2 and stimulated with Fsk/IBMX. **(D)** Representative confocal fluorescence images showing bleaching and recovery of an RIα condensate and diffuse region in a cell expressing RIα-GFP2 and Cα-mCherry. Scale bar, 10 µm. **(E)** Representative FRAP time-courses showing the fluorescence recovery of a photobleached RIα puncta in HEK293T cells expressing RIα-GFP2 alone (RIα-only puncta) or co-expressing Cα-mCherry plus either RIα-GFP2 (WT puncta) or RIα$^{A99S}$-GFP2 (A99S puncta). Fluorescence recovery of a photobleached cytosolic region from cells expressing RIα-GFP2 alone (Cytosol) is plotted as a reference. Data in **D** and **E** are representative of three independent experiments. **(F)** Representative image of *in vitro* colocalization of purified RIα (50 µM) and Cα (12.5 µM, 10% Cα-GFP2) with 10 µM cAMP. Scale bar, 10 µm. **(J)** AKAR4 basal Y/C emission ratio in RIα-KO HEK293T cells without (KO; n = 82 cells) or with expression of RIα-mRuby2 (RIα; n = 84 cells) or RIα$^{A99S}$-mRuby2 (A99S; n = 101 cells). Error bars indicate mean ± SD. ****P = $2.08 \times 10^{-9}$ (KO vs RIα) and ****P < $1\times10^{-15}$ (RIα vs A99S); Brown-Forsythe and Welch ANOVA followed by Dunnett's T3 multiple-comparisons test. **(G)** Normalized R/G emission ratio time-courses of RIα-KO HEK293T cells expressing FluoSTEP-AKAR with co-expression of RIα-FP11 (+RIα), RIα$^{A99S}$-FP11 (+A99S), or RIα-FP11 with Linker-mTagBFP2 (+RIα+Linker). **(H)** Basal AKAR4 Y/C emission ratio in RIα-KO HEK293T cells without (KO; n = 82 cells) or with expression of RIα-mRuby2 (RIα; n = 84 cells) or RIα$^{A99S}$-mRuby2 (A99S; n = 101 cells). Error bars indicate mean ± SD. ****P = $2.08\times10^{-9}$ (KO vs RIα) and ****P < $1\times10^{-15}$ (RIα vs A99S); Brown-Forsythe and Welch ANOVA followed by Dunnett's T3 multiple-comparisons test. **(I-J)** Normalized Y/C emission ratio time-courses of RIαKO HEK293T cells expressing **(I)** AKAR4 or **(J)** AKAR4(T/A) without (KO) or with RIα-mRuby2 (RIα) or RIα$^{A99S}$-mRuby2 (A99S) co-expression and stimulated with H89 (10 μM), followed by washout and addition of Fsk/IBMX. Curves plotted as means (solid lines) ± SD (shading). Data are from 3 independent experiments unless otherwise specified.

**Supplementary Table 1:** Residue clashes within D/D domain structure model containing L50R.

| Primary Residue | Atomic Coordinates | Clashing Residue | Atomic Coordinates | Overlap Distance (Å) |
|---|---|---|---|---|
| R50 | HH21 | L30’ | HD31 | 0.661 |
| I27 | HD12 | R50’ | HH22 | 0.604 |
| L36 | HD21 | R50 | CZ | 0.603 |
| R50 | HH12 | L31’ | HD21 | 0.597 |
| I27 | HG23 | R50’ | NH2 | 0.590 |
| R50 | HH22 | I27’ | HG23 | 0.584 |
| L36’ | HD22 | R50’ | CZ | 0.558 |
| R50 | HD3 | F54’ | CE1 | 0.558 |
| R50 | CG | R50 | HH11 | 0.529 |
| M47 | HA | R50 | HD2 | 0.503 |
| A48 | HA | G58 | OE1 | 0.486 |
| R50 | NH2 | I27’ | HD12 | 0.485 |
| I27 | HG23 | R50’ | HH22 | 0.484 |
| R50’ | HG3 | R50’ | HH11 | 0.469 |
| R50 | NH2 | L30’ | HD22 | 0.464 |
| R50 | CZ | L30’ | HD22 | 0.454 |
| L30 | HD13 | R50’ | NH2 | 0.416 |
| L38 | HD21 | R50 | NH1 | 0.401 |

**Supplementary Table 2:** Sequences of oligonucleotides used for plasmid construction (Related to STAR Methods)

| Type of constructs | Primer Name | Sequence (5’-3’) | Notes |
|---|---|---|---|
| RIα mutants | RIα_L50R_F | ATGGCATTCagaAGGGAATACTTTGAGAGGTTGGAG | Base changes are lower case; used with pcDNA3.1 GFP2-RIα, pcDNA3.1 mRuby2-RIα, and pRluc8-N3-hRIα constructs |
| | RIα_L50R_R | GTATTCCCTtctGAATGCCATGGGTCTCTCAGG | Base changes are lower case; used with pcDNA3.1 GFP2-RIα, pcDNA3.1 mRuby2-RIα, and pRluc8-N3-hRIα constructs |
| | RIα_Y122A_F | CCAAAAGATgcgAAGACAATGGCCGCTTTAGCCAAA | Base changes are lower case; used with pcDNA3.1 GFP2-RIα and pRluc8-N3-hRIα constructs |
| | RIα_Y122A_R | CATTGTCTTcgcATCTTTTGGTATAACCTTTCTAAC | Base changes are lower case; used with pcDNA3.1 GFP2-RIα and pRluc8-N3-hRIα constructs |
| | RIα_E202A_F | AGCTTTGGAgctCTTGCTTTGATTTATGGAACA | Base changes are lower case; used with pcDNA3.1 GFP2-RIα and pRluc8-N3-hRIα constructs |
| | RIα_E202A_R | GTTGGGGAAGGAGGGAGCTTTGGAgctCTTGCTTTG | Base changes are lower case; used with pcDNA3.1 GFP2-RIα and pRluc8-N3-hRIα constructs |
| | RIα_E326A_F | TATTTTGGTgccATTGCACTACTGATGAATCGTCCT | Base changes are lower case; used with pcDNA3.1 GFP2-RIα and pRluc8-N3-hRIα constructs |
| | RIα_E326A_R | TAGTGCAATggcACCAAAATAATCAGAAGGCCCCAA | Base changes are lower case; used with pcDNA3.1 GFP2-RIα |

| | | | |
|---|---|---|---|
| | | | and pRluc8-N3-hRIα constructs |
| | RIα_Nterm-linker-Ch2A_F | ACTgctACAgcaTCAgctgcagct gcaATTTCTCCTCCTCCAcct AACCCA gttGTTAAAGGTAGGAGGCG | Base changes are lower case; used with pcDNA3.1 GFP2-RIα constructs |
| | RIα_Nterm-linker-Ch2A_R | TGAtgcTGTagcAGTGCCTGC tgcCTGCAGATTCTGAATCT G tgcTGCCTCCTCCTTCTCCA ACCTCTCAA | Base changes are lower case; used with pcDNA3.1 GFP2-RIα constructs |
| | RIα_Cterm-linker-Ch2A_F | GcagctGCGGCATCCTCATGT TgcagctGTTATACCAgcagct TACAAGACAAATGGCCGC TT | Base changes are lower case; used with pcDNA3.1 GFP2-RIα constructs |
| | RIα_Cterm-linker-Ch2A_R | AACATAGGATGCCGCagctg cagcCGTGTAGACtgcAGCG CTGATAGCACCTCGTCGCC TC | Base changes are lower case; used with pcDNA3.1 GFP2-RIα constructs |
| | RIα_A99S_F | CGACGAGGTtCgATCAGCG CTGAGGTCTACACG | Base changes are lower case; used with pcDNA3.1 GFP2-RIα, pcDNA3.1 mRuby2-RIα, and pRluc8-N3-hRIα constructs |
| | RIα_A99S_R | AGGTAGGAGGCGACGAGG TtCgATCAGCGCT | Base changes are lower case; used with pcDNA3.1 GFP2-RIα, pcDNA3.1 mRuby2-RIα, and pcDNA3.1 FP11-RIα, pRluc8-N3-hRIα constructs |
| | RIα_A99E_F | GCGACGAGGTGaAATCAG CGCTGAGGTC | Base changes are lower case; used with pcDNA3.1 GFP2-RIα and pRluc8-N3-hRIα constructs |
| | RIα_A99E_R | CAGCGCTGATTtCACCTCG TCGCCTCC | Base changes are lower case; used with pcDNA3.1 GFP2-RIα and pRluc8-N3-hRIα constructs |
| | pRSET-RIα_A99S-F | GCGGCagtATTTCAGCGGAG GTT | Base changes are lower case; used with pRSET-RIα |

| | | | |
|---|---|---|---|
| | pRSET-RIα_A99S-R | CTGAAATactGCCGCGGCGACGAC | Base changes are lower case; used with pRSET-RIα |
| | pRSET-RIα_A99E-F | | Base changes are lower case; used with pRSET-RIα |
| | pRSET-RIα_A99E-R | | Base changes are lower case; used with pRSET-RIα |
| pET His6-SUMO-hPKAc-EGFP | hCa-linker-EGFP_F | GGCAAGGAGTTTTCTGAGTTTgGATCCCCACCGGTCGCC | Insert GFP C-terminus of human PKA-C |
| | hCa-linker-EGFP_R | GGCGACCGGTGGGGATCcAAACTCAGAAAACTCCTTGCCACAC | Insert GFP C-terminus of human PKA-C |
| | S76T_F | CTCGTGACCACCCTGAcCTACGGCGTG | Base changes are lower case; make GFP2 into EGFP |
| | S76T_R | CACGCCGTAGgTCAGGGTGGTCACGAG | Base changes are lower case; make GFP2 into EGFP |
| pcDNA3.1 AKB-mVenus-FKBP | smAKAP-AKB_R | ATATGGATACTCGCATGCCCATTGCTGCAA | Used to make both the AKB and AKB(L2P2)-FKBP constructs |
| | smAKAP-AKB_F | AGCGCCACCATGACTGTGATCTTGGAATAT | Used to make both the AKB and AKB(L2P2)-FKBP constructs |
| pcDNA3.1 GFP2-hCa | GFP2_F | AGACCCAAGCTGGCTAGCGTTTAAACTTAAGCTTGGGCCACCATGGTGAGCAAGGGCGAGGA | |
| | PKA_Ca_R | TTAAACGGGCCCTCTAGACTACTAAAACTCAGA | |
| | pcDNA3.1_backbone_F | CCAAGCTTAAGTTTAAACGCTAGCCAGCTTGGGTCTCC | |
| | pcDNA3.1_backbone_R | GTCTAGAGGGCCCGTTTAAACCCGCTGATCAGCCT | |
| pcDNA3.1 mRuby2-PKA-Ca | mRuby2-NTerm-F | GATCCGGGAGCATGGTGTCTAAGGGCGAAGAGCTGATC | |
| | PKAc-backbone-NTerm-R | CTTAGACACCATGCTCCCGGATCCAAACTCAGAAAACTCC | |
| | mRuby2-CTerm-R | TTAAACGGGCCCTCTAGACTATTACTTGTACAGCTCGTCCAT | |

| | | | |
|---|---|---|---|
| | PKAc-backbone-CTerm-F | TAGTCTAGAGGGCCCGTTTAA | |
| pEGFP-N1 smAKAP-FLAG-EGFP | smAKAP-NheI-F | GACgctagcACCATGGGCTGCATGAAATCAAAGC | NheI site is lower case |
| | smAKAP-FLAG-SalI-R-1 | GCgtcgacTGTTTATCGTCATCGTCTTTGTAGTC | SalI site is lower case. Underlined is the overlapping region. |
| | smAKAP-FLAG-SalI-R-2 | ATCGTCATCGTCTTTGTAGTCAGGCCCCTCACTCTCAATGTATGG | Underlined is the overlapping region. |
| pEGFP-N1 smAKAP(L2P2)-FLAG-EGFP | smAKAP-L2P2-F | GATCTTGGAATATGCACACCGCcCGTCTCAGGATATCccGTGTGATGCCTTGCAGCAATG | Base changes are lower case. |
| | smAKAP-L2P2-R | CTAGAACCTTATACGTGTGGCGgGCAGAGTCCTATAGggCACACTACGGAACGTCGTTAC | Base changes are lower case. |
| pcDNA3.1 mTagBFP-RIα(Linker) | pcBackbone_R | GGTGGCGGGTCTCCCTAT | Replicated from [Zhang et al 2020] |
| | RIα_AA62-113_R | GTGGCGACCGGTGGGGATCc GGATGCCGCATCTTCCTC | Replicated from [Zhang et al 2020] |
| | RIα _linker_F | gGATCCCCACCGGTCG | Replicated from [Zhang et al 2020] |
| | pcBackbone_mTagBFP2_R | GCTGGATATCTGCAGAATTC TTAATTAAGCTTGTGCCCCAGT | Replicated from [Zhang et al 2020] |
| pcDNA3.1 FP11-RIα | RIα-FP11_F | GTGTCACTGTCTGTC GGAGGAACAGGAGGTT | |
| | RIα-FP11_R | CTGCAGAATTC TCATGTAATCCCAGC | |
| | FP11_F | GGAGGAACAGGAGGTTCACGTGACCACATGGTCCTTCATGAGTATGTAAATGCTGCTGGGATTACATGA | |
| | RIα_N_R | GACAGACAGTGACACAAAACTGTT | |
| | pcBackbone_F | GAATTCTGCAGATATCCAGCACAGTGG | |
